# Klf9 is a key feedforward regulator of the transcriptomic response to glucocorticoid receptor activity

**DOI:** 10.1101/863555

**Authors:** Ian Gans, Ellen I. Hartig, Shusen Zhu, Andrea R. Tilden, Lucie N. Hutchins, Nathaniel J. Maki, Joel H. Graber, James A. Coffman

**Affiliations:** MDI Biological Laboratory, Salisbury Cove, Maine, USA; University of Maine Graduate School of Biomedical Sciences and Engineering, Orono, Maine, USA; Colby College, Waterville, Maine, USA

**Keywords:** Glucocorticoid receptor, Krüppel-like Factor 9, Cortisol, Zebrafish, Transcriptomics, RNA-seq, CRISPR

## Abstract

The zebrafish has recently emerged as a model system for investigating the developmental roles of glucocorticoid signaling and the mechanisms underlying glucocorticoid-induced developmental programming. To assess the role of the Glucocorticoid Receptor (GR) in such programming, we used CRISPR-Cas9 to produce a new frameshift mutation, GR^369-^, which eliminates all potential in-frame initiation codons upstream of the DNA binding domain. Using RNA-seq to ask how this mutation affects the larval transcriptome under both normal conditions and with chronic cortisol treatment, we find that GR mediates most of the effects of the treatment, and paradoxically, that the transcriptome of cortisol-treated larvae is more like that of larvae lacking a GR than that of larvae with a GR, suggesting that the cortisol-treated larvae develop GR resistance. The one transcriptional regulator that was both underexpressed in GR^369-^ larvae and consistently overexpressed in cortisol-treated larvae was *klf9*. We therefore used CRISPR-Cas9-mediated mutation of *klf9* and RNA-seq to assess Klf9-dependent gene expression in both normal and cortisol-treated larvae. Our results indicate that Klf9 contributes significantly to the transcriptomic response to chronic cortisol exposure, mediating the upregulation of proinflammatory genes that we reported previously.

## Introduction

The vertebrate hypothalamus-pituitary-adrenal (HPA) axis orchestrates physiological, behavioral, and metabolic adjustments required for homeostasis, by dynamically regulating production and secretion of adrenal steroids known as glucocorticoids. In humans the primary glucocorticoid is cortisol, the biological activity of which is mediated by two regulatory proteins in the nuclear receptor family, the ubiquitous glucocorticoid receptor (GR) and the more tissue-restricted mineralocorticoid receptor (MR). The GR binds cortisol less avidly than the MR and is thus more dynamically regulated over the normal physiological range of cortisol fluctuations^1,2^. The GR and MR function both as transcription factors and as non-nuclear signaling proteins, including in the central nervous system where both proteins are highly expressed^1-5^. Given that the GR is more widely expressed and more dynamically regulated by cortisol, it is generally thought to be the principal mediator of cortisol-induced genomic responses to circadian rhythms and acute stress^5^.

A GR target gene that was recently shown to mediate circadian regulation of cell proliferation in human epidermis is *KLF9*^6^, which encodes a ubiquitously expressed^7^ member of the krüppel-like family of zinc finger transcription factors. Klf9 is a repressor important for neurogenesis and control of neural plasticity^8-11^, and is a central player in the GR-responsive gene regulatory network in macrophages^12^, where it functions to repress another GR target gene, *klf2*. In this context the GR, *klf9*, and *klf2* constitute a regulatory circuit known as an ‘incoherent type-1 feed-forward loop’ (i1-FFL)^12^, a network motif commonly used to generate pulsatile dynamics and accelerate responses^13-15^. GR signaling often exhibits incoherent feed-forward regulatory logic^16-18^. In epidermis many of the genes repressed by Klf9 are also activated by the GR^6^. It was recently shown that *klf9* is upregulated in mouse hippocampus in response to the onset of chronic stress, and blocking that upregulation prevents chronic stress-induced pathologies^19^. In the same study it was also shown that hippocampuses of women with major depression were found to overexpress *KLF9*^19^. Finally, it was recently discovered that Dexamethasone-induced upregulation of Klf9 promotes hepatic gluconeogenesis and hyperglycemia in mice^20^. Thus, *klf9* is a GR target gene that dynamically controls GC signaling, and its overexpression is implicated in stress-induced pathology.

The zebrafish has recently emerged as a model system well-suited to investigating the developmental functions of glucocorticoid signaling and mechanisms underlying developmental programming in response to chronic glucocorticoid exposure such as occurs with chronic early life stress^21-25^. As in humans, the endogenous glucocorticoid in zebrafish is cortisol, which is produced by the interrenal gland, the functional equivalent of the mammalian adrenal. Thus, the zebrafish homolog of the HPA axis is the Hypothalamus-Pituitary-Interrenal (HPI) axis. Eight different zebrafish GR mutants have been described in the literature to date^26-29^. The first of these, *gr*^*s357*^, is a missense mutation that substitutes a cysteine for an arginine in the DNA binding domain, abolishing GR DNA binding activity^26^. The other mutations, all produced by targeted mutagenesis using CRISPR-Cas9 or TALEN technology, consist of frameshift indels in exons 2 and 5^27-29^, which respectively encode domains N- and C-terminal to the DNA binding domain (see Fig. 1). All of the zebrafish GR mutants manifest behavioral defects but are viable as homozygotes.

**Figure 1.**
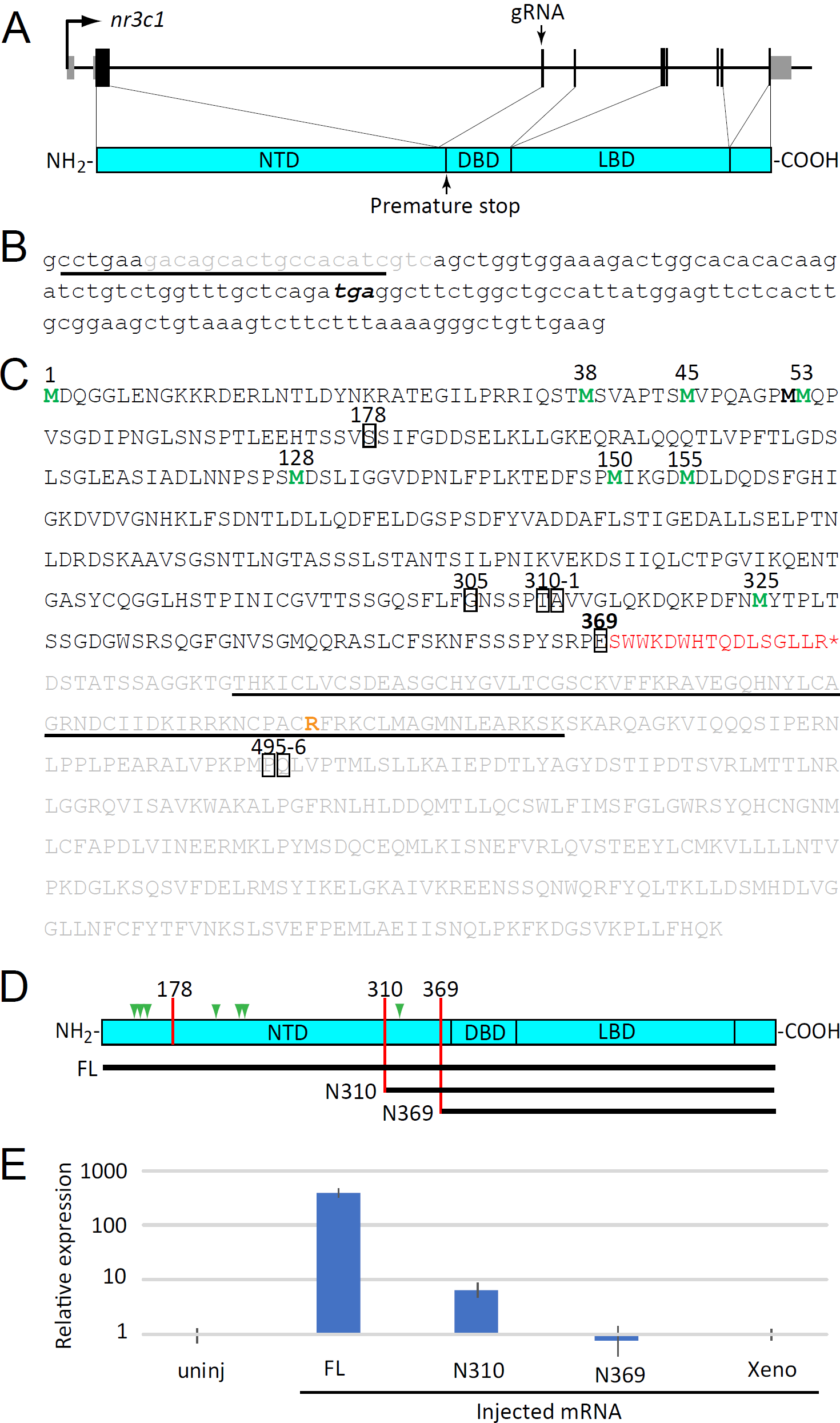
A new CRISPR-Cas9-induced mutation of the zebrafish GR. (A) Schematic of the *nr3c1* gene and encoded GR showing location of the targeted sequence and resulting premature stop codon with respect to the protein domains: N-terminal domain (NTD), DNA binding domain (DBD), and ligand binding domain (LBD). (B) Nucleotide sequence of nr3c1 exon 3, showing sequence targeted by the gRNA (underlined), the 20 base deletion resulting from injection of that gRNA and Cas9 mRNA (gray), and the resulting premature stop codon (bold italic). (C) Predicted amino acid sequence of the full-length zebrafish GR, plus the extra amino acids introduced by the frameshift shown in (B) (red font). The sequence eliminated by the premature stop is shown in gray, with the DBD underlined. Methionines corresponding to potential alternative initiation sites are shown in green. The positions of all reported frameshift mutations (including that reported here, E369) are boxed. The arginine that is changed to a cysteine in the *gr*^*s357*^ mutation is shown in orange. (D) Schematic of the GR showing locations of the two previously reported exon 2 frameshift mutations resulting in truncations at amino acids 178 and 310, and the one in exon 3 reported here truncating at amino acid 369, and residues included in N310 and N369 mRNA constructs produced for microinjection. (E) Relative expression of *fkbp5* in 6 hour homozyogous GR^369-^ embryos, either uninjected or injected with mRNA encoding full-length GR (FL), N-terminally truncated isoforms N310 and N369, or a nonspecific control mRNA encoding *Xenopus* elongation factor 1α (Xeno).

We showed previously that zebrafish larvae treated chronically with 1 μM cortisol upregulate proinflammatory genes and give rise to adults that maintain chronically elevated cortisol and dysregulated expression of those same genes^23^. Thus in zebrafish, chronic cortisol exposure during early development results in persistent, long-term dysregulation of the HPI axis and gene expression downstream thereof, a ‘developmental programming’ effect similar to what has been observed with early life stress in mammals^23,25^. To ask whether and to what extent that developmental programming depends on the GR, we used CRISPR-Cas9 technology to mutate the GR*-*encoding gene *nr3c1*. Here we report a new loss-of-function mutation, consisting of a deletion in *nr3c1* exon 3 that results in a frameshift and premature stop codon immediately upstream of the DNA binding domain which eliminates GR activity as a transcriptional activator. Using this mutant line, we performed an RNA-seq experiment to identify genes regulated by the GR and to parse GR-dependent and GR-independent effects of chronic cortisol treatment on larval gene expression. We found that most of the transcriptomic effects of the chronic cortisol treatment are mediated by the GR and identified *klf9* as the one transcriptional regulatory gene consistently upregulated by the GR under both normal conditions and in response to chronic cortisol treatment. We therefore used CRISPR-Cas9 to mutate *klf9* and RNA-seq to interrogate its role with respect to the transcriptomic response to chronic cortisol treatment. Our results indicate that *klf9* mediates the cortisol-induced upregulation of pro-inflammatory genes that we previously reported^23^, and suggest that *klf9* is a core feedforward regulator of the transcriptional response to glucocorticoid signaling.

## Results

### A frameshift deletion introduced into exon 3 of zebrafish *nr3c1* eliminates GR transcriptional function

To mutate the GR, we employed CRISPR-Cas9 using a guide RNA (gRNA) that targets *nr3c1* exon 3 (the second coding exon; Fig. 1A). This resulted in a 20 base deletion and frameshift that introduces sixteen new amino acids after E369, followed by a premature translational stop immediately upstream of the DNA binding domain (Fig. 1B, C), a mutation hereafter referred to as GR^369-^. F0 males with GR^369-^ in the germline were identified and outbred to wild-type (AB) females, and siblings of that cross were mated to generate F2s, from which homozygous males and females were identified. Homozygous GR^369-^ males crossed with wild-type females produced viable embryos, whereas most homozygous GR^369-^ females crossed with either mutant or wild-type males produced embryos that all died within 24 hours of fertilization. However, one produced 10 viable offspring that were raised to adulthood, indicating that maternal GR is not essential for early development. When inbred these F3 fish produced no viable offspring. However, females homozygous for GR^369-^ following another generation of outcrossing produced variable numbers of viable offspring. These results suggest that loss of GR function may compromise egg quality, albeit in a context-dependent way.

The zebrafish *nr3c1* transcript has multiple AUG codons that could potentially initiate translation to allow production of alternative N-terminal isoforms from a single mRNA (Fig. 1C, D; Fig. S1), as occurs with human *nr3c1*^30^ and some other transcription factors (e.g.^31^). One potential in-frame initiation codon upstream of the DNA binding domain lies downstream of all previously reported frameshift mutations in exon 2^27-29^ (Fig. 1 C,D), suggesting that *nr3c1* mRNA transcribed in those mutants might still allow translation of an N-terminally truncated GR isoform with an intact DNA binding domain. To assess the transcriptional activity of different N-terminal isoforms we injected homozygous GR^369-^ zygotes with mRNA encoding full-length GR (FL) or GRs lacking the first 310 or 369 amino acids of the full-length GR (N310 and N369, Fig. 1D). The N310 mRNA corresponds to the shortest translation initiation variant with a DNA binding domain potentially available to the previously described exon 2 mutants^27-29^, whereas the N369 corresponds to what is available in GR^369-^ mutants. The embryos were collected at 6 hours post fertilization (hpf) and quantitative reverse transcription and polymerase chain reaction (qRT-PCR) was used to measure the expression of the GR-target gene *fkbp5*. Unlike uninjected embryos or embryos injected with N369 or a *Xenopus* control mRNA, embryos injected with N310 mRNA upregulated *fkbp5*, indicating that N310 retains activity as a transcription factor (Fig. 1E). Zygotes injected with mRNA encoding full length GR showed ∼50-fold stronger upregulation of *fkbp5* than those injected with N310 (Fig. 1E). We conclude that the previously described *nr3c1* exon 2 frameshift mutations^27-29^ strongly abrogate but do not abolish the potential to translate a GR with transcriptional function, whereas GR^369-^ eliminates potential for translating a transcriptionally active GR.

### RNA-seq shows that the transcriptomic response to chronic cortisol treatment is largely GR-dependent but converges with that caused by loss of GR function

To identify GR-dependent zygotic gene expression and assess the extent to which the GR contributes to the transcriptional effects of chronic cortisol exposure^23^, we used a visual background adaptation (VBA) assay to identify homozygous GR^369-^ larvae from a heterozygous cross. This assay makes use of the fact that larval melanophores mount a camouflaging response to changes in background, dispersing when larvae are transferred from the dark to a light background^26^. This response is mediated by the GR, so larvae lacking a functional GR can be identified as those in which melanophores fail to respond, remaining clustered in a dark patch after transfer to a light background. At four days post fertilization (dpf) we used the VBA screen to separate homozygous GR^369-^ (VBA-) larvae from their heterozygous and wild type (VBA+) siblings developed under normal (vehicle-treated) conditions or in medium supplemented with 1 μM cortisol (Fig. 2A). In support of the screen’s validity, the number of VBA-larvae was ∼1/4 of the total number of larvae, the expected Mendelian ratio for homozygous mutants. The larvae developed for another day under the same conditions (vehicle- or cortisol-treated), at which time four biological replicates of each sample were collected for RNA-seq (Fig. 2A).

**Figure 2.**
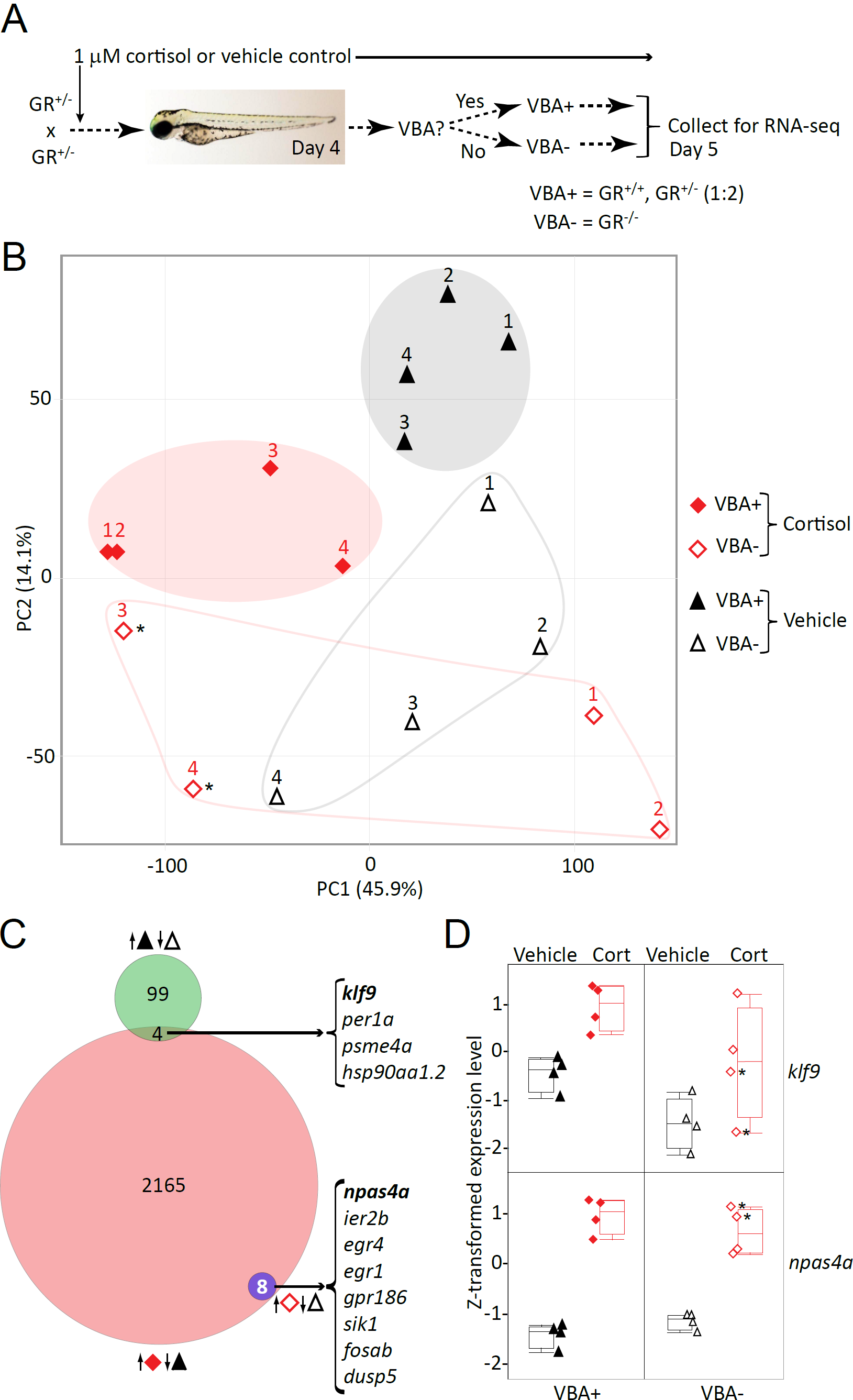
RNA-seq identifies GR-dependent and GR-independent genes and effects of chronic cortisol treatment. (A) Diagram of experimental design using Visual Background Adaptation (VBA) selection. (B) Principal component plot of the RNA-seq data, showing the location of each replicate sample with respect to the first two principal components. Cortisol-treated samples are shown in red, vehicle-treated controls in black. VBA+ samples (containing at least one functional GR allele) are filled shapes, VBA-samples are empty shapes. (C) Venn diagram showing numbers of genes upregulated in each of the indicated comparisons (keyed as in (A)): VBA+ vs. VBA-(vehicle-treated controls); cortisol-treated vs. vehicle-treated VBA+; and cortisol-treated vs. vehicle treated VBA-. Upregulated genes in common respectively between the first two and the last two comparisons are listed on the right. (D) Box plots of Z-transformed expression levels of *klf9* and *npas4a* obtained from the RNA-seq data.

A principal component (PC) analysis of the RNA-seq data indicated that 60% of the variance in gene expression among samples is captured in the first two PCs, which respectively correlate with the two treatments (chronic cortisol and absence of a GR, respectively accounting for 46% and 14% of the variance; Fig. 2B). The eight VBA+ (i.e. mixed wild type and heterozygous mutant) samples cluster according to whether they were treated with cortisol, with the two clusters (cortisol-treated and vehicle-treated controls) segregating toward opposite poles of PC1. This is not the case in the VBA- (i.e. homozygous GR^369-^) fish, the four cortisol-treated replicates of which are widely dispersed along PC1. This indicates that the global effect of the cortisol treatment captured in PC1 is GR-dependent and suggests as well that gene expression is less constrained overall in larvae lacking a GR. PC2 correlates with the presence and absence of a GR (VBA+ vs. VBA-respectively). Interestingly, along PC2 chronic cortisol treatment displaces the VBA+ transcriptome toward the VBA-pole, suggesting that the cortisol-treated fish adapt to the exposure by developing GR resistance.

The regulatory roles of the GR were further assessed by analyzing differential gene expression (DGE) between pairs of treatments, using an adjusted *p*-value (false-discovery) threshold of 0.05 as the criterion for differential expression. Comparison of VBA+ and VBA-larvae identified 405 genes affected by loss of the GR in 5-day larvae, 103 of which are underexpressed (Fig. 2C) and 302 of which are overexpressed in VBA-larvae compared to their VBA+ counterparts (Fig. S2A). Gene ontology (GO) analysis shows that the underexpressed genes (i.e. genes normally upregulated by the GR) are involved in sugar metabolism and response to heat, whereas the overexpressed genes (i.e. genes that are normally downregulated via the GR) are involved in basement membrane organization, epidermis development, cell adhesion, locomotion, and growth (Figs. S3 and S4; Table S1). A DGE analysis comparing cortisol-treated VBA+ fish to their vehicle-treated VBA+ counterparts showed that in cortisol-treated larvae with a functional GR, 4,298 genes were differentially expressed (Fig. S2B), 2,177 of which were upregulated (Fig. 2C) and 2,121 of which were downregulated. GO enrichment analysis of the upregulated genes identified biological processes related to nervous system development and function as well as cell adhesion, locomotion, and growth, whereas the downregulated genes were largely involved in protein synthesis and metabolism (Figs. S5, S6; Table S2). Interestingly, the transcriptome of cortisol-treated VBA+ larvae overlapped more with that of vehicle-treated VBA-larvae than with that of vehicle treated VBA+ larvae (Fig. S2C), and accordingly, many of the biological processes affected by the absence of GR function in VBA-larvae were similarly affected by the chronic cortisol-treatment in VBA+ larvae (Fig. S7). This again suggests that the latter larvae develop a GR resistant phenotype.

We reasoned that some of the effects of chronic cortisol exposure might stem from GR-induced upregulation of a GR target gene that functions as a feedforward transcriptional regulator of GR signaling. Such a gene should be both underexpressed in VBA-larvae and upregulated in VBA+ larvae in response to chronic cortisol (i.e., opposite of the predominant trend noted above). Of the 2,177 genes upregulated in cortisol-treated VBA+ larvae only four were basally underexpressed in VBA-larvae (Fig. 2C), two of which encode transcription factors: *klf9* and *per1a*. Of these, only *klf9* was also found to be significantly upregulated in our previous RNA-seq analysis of the effects of chronic cortisol treatment in wild-type fish (Fig. S2C), being one of the most highly upregulated transcription factors^23^. A plot comparing *klf9* expression in each of the conditions reveals that the GR contributes to both its normal developmental expression and its upregulation in response to chronic cortisol treatment (Fig. 2D). However, the plot also suggests that cortisol affects *klf9* in a GR-independent fashion, albeit more variably, as indicated by the range of expression levels in the cortisol-treated VBA-samples shown in Fig. 2D, which correlate with the spread of the cortisol-treated VBA-samples along PC1 shown in Fig. 2B.

In contrast to the situation in VBA+ larvae, only 8 genes were differentially expressed in cortisol-treated VBA-larvae compared to their vehicle-treated VBA-siblings, all of them upregulated (Fig. 2C, Fig. S2D). The genes included the immediate early genes (IEGs) *npas4a, egr1, egr4, fosab*, and *ier2b* (Fig. 2C, Fig. S8). The GR-independence of their cortisol-induced upregulation is clearly seen in a plot of the expression levels of the most highly upregulated gene of this set, *npas4a* (Fig. 2D), a neuronal IEG that along with the other IEGs was also found to be upregulated in our previous RNA-seq analysis of cortisol-treated larvae (Fig. S2D)^23^. This indicates that the GR mediates nearly all the transcriptomic effects of chronically elevated cortisol, except for a small subset that appears to relate to neuronal activity.

### A frameshift deletion introduced into exon 1 of zebrafish *klf9* eliminates the DNA binding domain and significantly reduces expression of the mature transcript

The fact that *klf9* was the transcriptional regulatory gene most consistently found to be upregulated by chronic cortisol exposure in a GR-dependent way suggested that it may contribute to the transcriptomic effects of the exposure. To test this, we mutated *klf9* using CRISPR-Cas9 with a gRNA that targets exon 1 (Fig. 3A, B). This resulted in a frameshift mutation upstream of the DNA binding domain (Fig 3A, B), producing a transcript encoding a truncated protein predicted to lack function as a transcription factor. *Klf9* loss-of-function mutations are viable in mice^32^ and similarly, the *klf9* mutant fish were viable and fertile when bred to homozygosity, although mutant embryos survive at a lower rate than wild type (data not shown). To ask how the mutation affects *klf9* expression we used quantitative reverse transcription and polymerase chain reaction (qRT-PCR) to compare *klf9* transcript levels in wild type and *klf9* homozygous mutant (hereafter referred to as *klf9*^-/-^) larvae under both normal conditions and in response to chronic cortisol treatment. This provided further confirmation that chronic cortisol treatment leads to upregulation of *klf9* and revealed that *klf9* mRNA levels are significantly reduced in the *klf9*^-/-^ larvae, probably due to nonsense-mediated decay triggered by the premature stop codon (Fig. 3C). In support of this possibility, there was no significant effect on *klf9* pre-mRNA levels, measured by qRT-PCR of the intron (Fig. 3C). We conclude from these experiments that the frameshift mutation introduced into *klf9* exon 1 abrogates Klf9 function.

**Figure 3.**
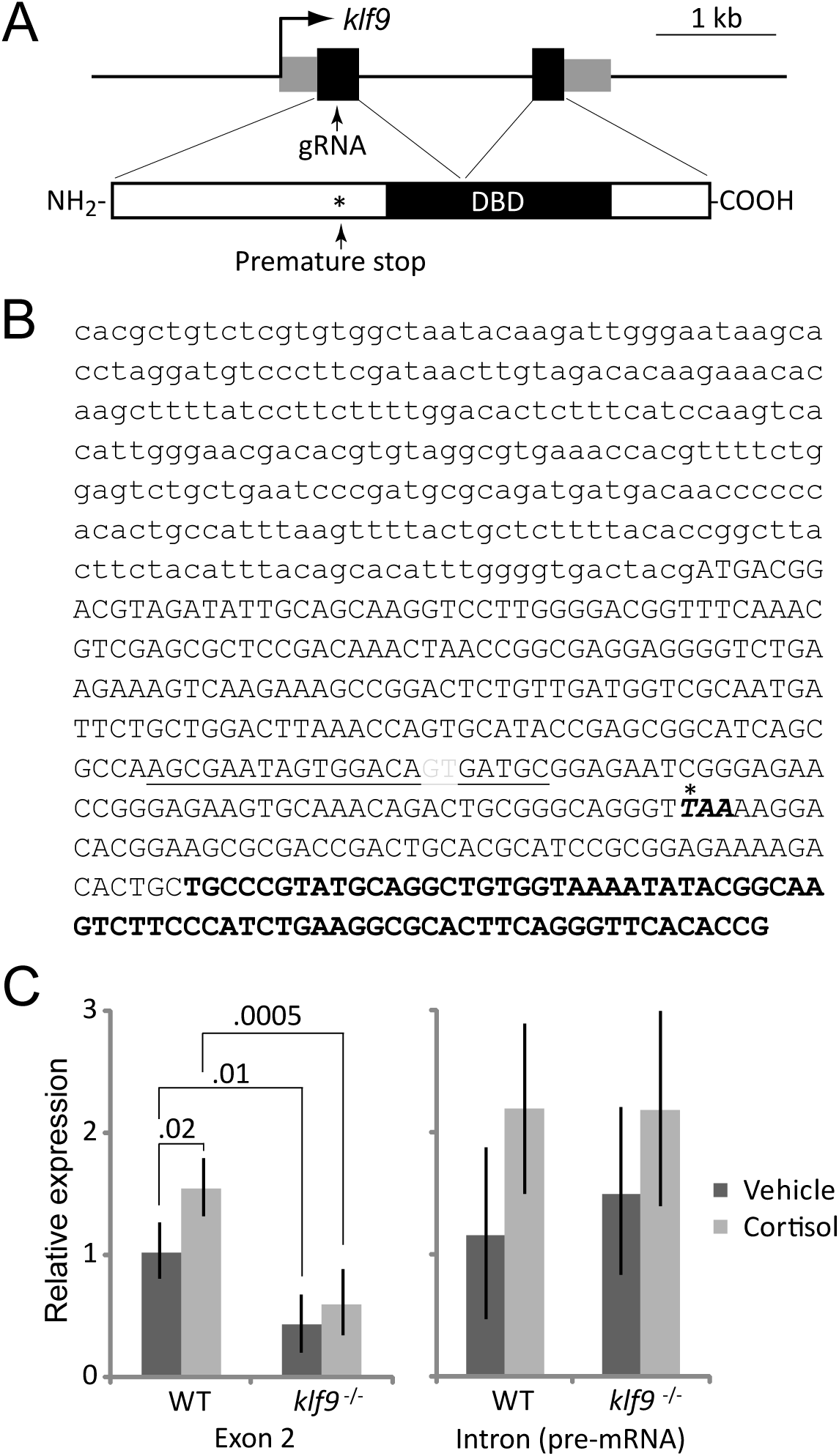
CRISPR-Cas9-mediated disruption of zebrafish *klf9*. (A) Schematic of the *klf9* locus and encoded protein showing location of the gRNA target in exon 1, which introduces a premature stop codon upstream of the DNA binding domain. (B) Sequence of *klf9* exon 1, with 5’ untranslated region in small case and coding sequence in upper case. The location of the 2 base pair deletion generated by CRISPR is shown in gray font; the gRNA target sequence is underlined; and the premature stop codon introduced by frameshift is noted with an asterisk. (C) Effect of chronic cortisol treatment and the frameshift mutation on relative levels of *klf9* mRNA and pre-mRNA, measured by qRT-PCR. The results show the averages and SEM of three biological replicates; significance values were obtained by ANOVA.

### RNA-seq shows that *klf9* regulates immune gene expression and is required for the pro-inflammatory transcriptomic effects of chronic cortisol treatment

To identify Klf9 target genes and ask whether Klf9 contributes to the transcriptomic response to chronic cortisol treatment we used RNA-seq to query gene expression in 5-day old wild type and *klf9*^-/-^ mutant larvae from sibling parents, developed both normally and in the presence of 1 μM cortisol. Samples were collected at the same time on day 5 post-fertilization as in the previous GR knockout experiment and processed similarly. However, PC analysis revealed the largest source of variance in this experiment was not due to genotype or treatment, but rather the order in which the samples were collected and processed (Fig. S9A). GO analysis of genes correlated with the first principal component showed that the later samples (replicates 3 and 4) had increased synaptic signaling and decreased translation (Table S3), suggestive of a physiological stress response (e.g. to the stress of capture). However, a further confound is that PC1 also correlates with preparation of the RNA in two batches on separate days, suggesting that it may also reflect technical variance in sample preparation (see Materials and Methods). Reassuringly, after normalizing for this variation the samples segregate along two principal components representing genotype and treatment (Fig. S9B). DGE analysis using a false-discovery rate of .05 identified 77 genes affected by loss of Klf9 function in vehicle-treated larvae, 33 of which were upregulated and 44 of which were downregulated. Gene ontology term enrichment analysis shows that the upregulated genes are involved in chromatin assembly and epigenetic regulation, as well as activation of immune response, immune system process, ATP metabolism, carbohydrate catabolism and response to stimulus (Fig. S10, Table S4), whereas the downregulated genes are involved in sterol biosynthesis, reproductive structure development, response to glucocorticoid, cell activation and epigenetic regulation (Fig. S11, Table S4).

To ask how loss of klf9 function affects the transcriptomic response to chronic cortisol exposure, we used hierarchical clustering to identify genes that were differentially expressed in cortisol-treated wild-type larvae but not cortisol-treated *klf9*^-/-^ larvae. This identified 228 genes upregulated by the cortisol-treatment in a klf9-dependent way (Fig 4A, B and Fig. S12), about a fifth (43) of which were also shown by our previous study^23^ to be significantly upregulated by chronic cortisol exposure. Examples of the latter include *irg1l* and *marco* (Fig. 4C and Fig. S13). Gene ontology term enrichment analysis of the 228 genes upregulated by chronic cortisol treatment in wild-type but not *klf9*^-/-^ embryos revealed significant enrichment for genes involved in defense and immunity (Fig. 4D, Table S5), the same biological processes that we previously found to be the most strongly affected by the chronic cortisol treatment^23^. These results indicate that Klf9 contributes in a significant way to the pro-inflammatory gene expression induced by chronic cortisol exposure.

**Figure 4.**
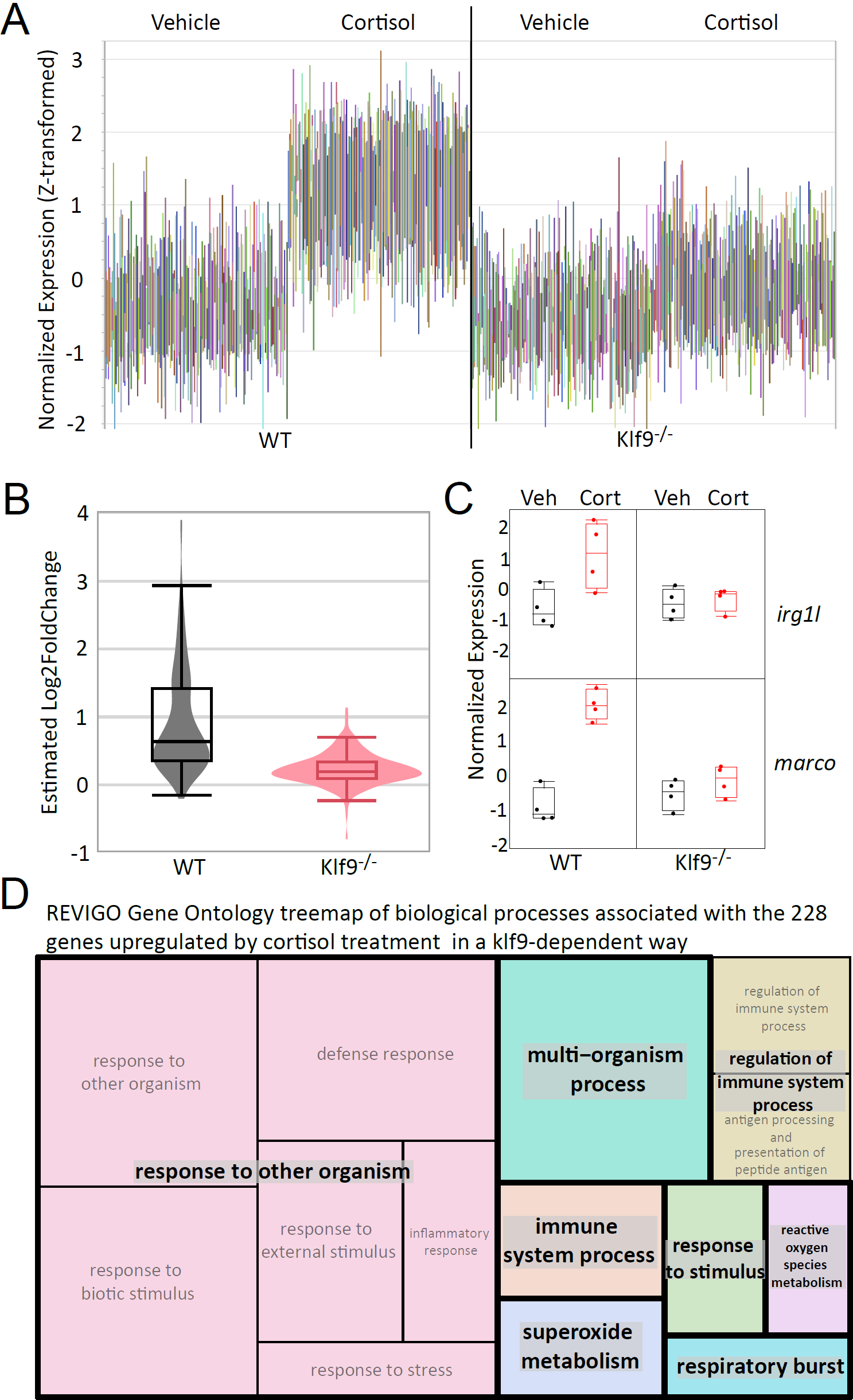
RNA-seq identifies *klf9*-dependent transcriptomic effects of chronic cortisol exposure. (A) Box plots of Z-transformed normalized expression levels of genes upregulated by chronic cortisol treatment in wild type larvae but not in *klf9*^-/-^ larvae. (B) Violin and box plots comparing estimated fold change of the 228 genes identified in (A); a nonparametric comparison using the Wilcoxon Method shows that the difference is statistically significant (p < .0001; Table S11). (C) Examples of two of the genes upregulated by chronic cortisol in a klf9-dependent fashion that were also identified in our previous analysis. (D) REVIGO treemap of gene ontology biological process terms found by GOrilla to be enriched in the set of genes upregulated by chronic cortisol in a *klf9*-dependent way.

### Genes found to be consistently upregulated by chronic cortisol treatment in multiple RNA-seq experiments depend on *klf9* for that upregulation

As a final analysis we assessed the overlap between the transcriptomic effects of chronic cortisol treatment across all our RNA-seq experiments with wild-type and VBA+ (mixed wild-type and heterozygous GR^369-^) larvae, including the experiment published previously^23^. To eliminate any technical artifacts emanating from the use of different parameters in the different analyses the sequence reads from all the experiments were reanalyzed from scratch using a common pipeline (see Materials and Methods). A PC analysis of the variance across all experiments produced several interesting observations. First, it showed that the effect of chronic cortisol treatment across all experiments was subtle, found only in PCs 4, 5, and 6 accounting for 14.3% of the total variance. PC5 captures a cortisol-treatment effect common to all three experiments (accounting for 2.77% of the total variance) and when plotted against PC4 clearly shows segregation of the cortisol-treated and control (vehicle-treated) samples (Fig. 5A). Gratifyingly, GO analysis of a single gene list ranked by upregulation along PC5 showed the same biological response to chronic cortisol as that which we reported previously^23^, i.e. upregulation of processes related to defense, inflammation and immunity (Fig. 5B, Table S6). The fourth principal component, accounting for 8.95% of the total variance, shows a cortisol treatment effect in the two experiments reported here, but not in that which we reported previously^23^ (Fig. 5A and Fig. S14). This suggests that PC4 represents cortisol-treatment effects that are dependent on the circadian light-dark cycle under which embryos in this study developed, which was absent in our previous study in which embryos developed in the dark^23^ (see below, Discussion). The sixth PC (2.57%) is the complement of PC4, suggesting cortisol-induced effects that are only apparent in the larvae that were developed in the dark (Fig. S14).

**Figure 5.**
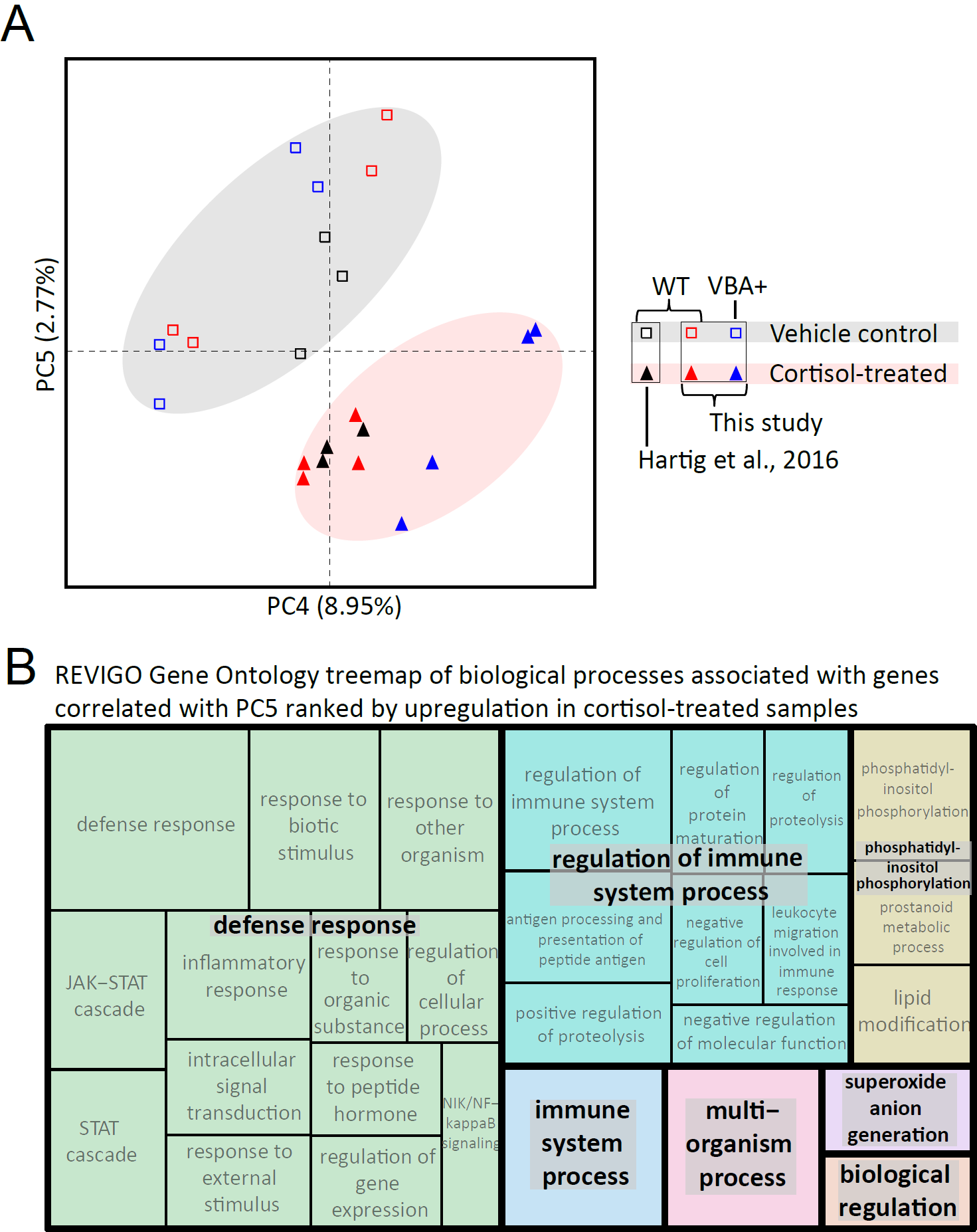
Combined analysis of three different RNA-seq datasets examining the transcriptomic effects of chronic cortisol exposure. (A) PC plot showing the location of each wild-type or VBA+ sample from all three RNA-seq experiments with respect to PCs 4 and 5. (B) REVIGO treemap of GO biological process terms found by GOrilla to be associated with PC5 ranked by upregulation in response to chronic cortisol treatment.

The spread of the cortisol treatment effects across multiple principal components likely reflects differing biological responses to the treatment among different experiments. That the effects are somewhat different under total dark compared with light-dark cycles is not surprising given what is known about the interplay of GC signaling and the circadian clock^33-36^. Indeed GO analysis of PC2, which segregates our previously published experiment^23^ from the two reported here (accounting for 18.5% of the variance), clearly shows that effects on circadian and light-responsive gene expression (Table S7). Additional sources of variance are less clear. The first PC, which segregates the wild-type samples in the last (klf9^-/-^ vs. wild-type) experiment from the wild-type samples in our previous experiment as well as the VBA+ samples described here (accounting for 37.6% of the variance, Fig. S14), is heavily loaded with genes involved in synaptic signaling and neurogenesis (Table S8) suggestive of differences in neurodevelopment and/or responsiveness to stress in those samples. The third PC (14.2% of the variance) segregates replicates 1 and 2 of the Klf9 experiment from all other samples, possibly reflecting the batch effect in sample preparation noted above.

Unsurprisingly given the large amount of gene expression variance across experimental samples unrelated to the cortisol treatment (i.e. noise), only 12 genes were found to be consistently upregulated by the cortisol treatment in all the experiments with statistical significance (adjusted p < 0.05), whereas none were consistently downregulated in all experiments (Fig. 6A). The consistently upregulated set included *klf9*, the only gene in that set that encodes a transcription factor. We reasoned that many more genes are affected by the treatment, albeit not consistently with statistical significance owing to the abovementioned noise (see Discussion). This was borne out by plotting the estimated fold-change of the 149 genes that were upregulated by cortisol-treatment in at least two of the three experiments (Fig. 6B), which also revealed that the upregulation of those genes is klf9-dependent. Furthermore, this plot shows that the cortisol treatment effect is stronger in the wild-type larvae than in the VBA+ (1:2 wild-type:heterozygous GR^+/-369-^), suggesting haploinsufficiency of the GR for some of the effect. Query of this set for enrichment of GO biological process terms indicated chronic cortisol exposure upregulates genes associated with response to organic substance (including response to stress, defense response, response to external biotic stimulus, and inflammatory response to wounding), similar to what we reported previously, and gluconeogenesis (Fig. 6C, Table S9).

**Figure 6.**
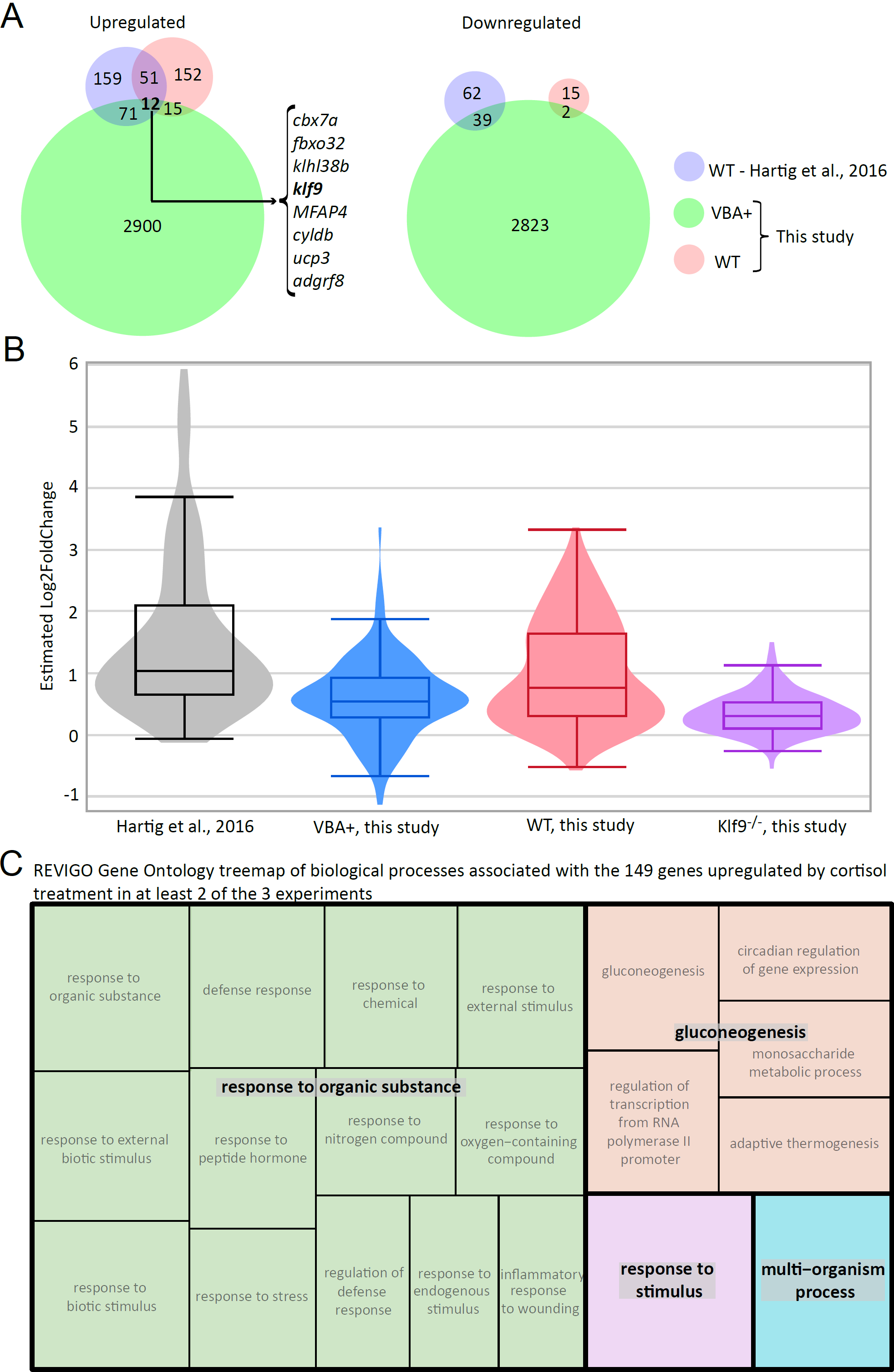
Identification and analysis of a common set of genes upregulated by chronic cortisol treatment in multiple RNA-seq experiments. (A) Venn diagrams showing numbers of upregulated and downregulated genes following reanalysis of the data from all three experiments and their overlap. Only 8 of the 12 genes in common between all experiments were annotated with gene names (listed). (B) Violin and box plots of estimated fold change of the 149 genes that are upregulated in at least 2 of the three experiments shown in (A); the differences between each experiment are statistically significant (Table S12). (C) REVIGO treemap of GO biological process terms found by GOrilla to be associated with the 149 genes upregulated by chronic cortisol treatment in at least two out of the three experiments.

Finally, we used HOMER motif enrichment analysis^37^ to ask what transcription factor binding sites are enriched in flanking regions of the set of 149 genes upregulated by cortisol-treatment in at least two of the three experiments. The resulting set of motifs included sites for the various krüppel-like factors as well as the GR (Table S10). The most significantly enriched motif, the Klf14 binding motif RGKGGGCGKGGC, matches the Klf9 consensus motif in the JASPAR database^38,39^ and would be expected to bind Klf9, which is in the same KLF subfamily as Klf14^40,41^. In contrast, these sites were not found to be enriched in the set of 228 genes identified in Figure 4 (which was instead enriched for several immunoregulatory transcription factors; Table S10; note that this set has 44 genes in common with the 149), suggesting that that larger set includes more indirect targets of feedforward regulation downstream of klf9. Consistent with this, the latter set of motifs includes sites for two immunoregulatory genes, *irf1* and *stat4* that are both consistently upregulated by chronic cortisol treatment (i.e. in the common list of 149 genes) and dependent on klf9 for that upregulation (i.e. in the list of 228 Klf9-dependent genes). It is also worth noting however that the set of 149 genes was identified by the more stringent criterion of being upregulated in multiple experiments and is therefore likely to have fewer false positives. Altogether these results underscore the conclusion that *klf9* is a feedforward regulator of GR signaling that mediates the pro-inflammatory transcriptomic response to chronic cortisol exposure, likely via a combination of direct effects on Klf9 regulatory targets and indirect effects of transcriptional regulatory genes downstream thereof.

## Discussion

The zebrafish has recently emerged as a model system for investigating the developmental functions of the GR and its role in glucocorticoid-induced developmental programming. Several previously published studies have characterized loss-of-function mutations of the zebrafish GR, including four frameshifting indels that introduce a premature stop codon in exon 2^27-29^ (Fig. 1C). Unlike those mutations, our frameshift deletion (GR^369-^) lies within exon 3 (Fig. 1A, B), which eliminates all possible in-frame initiation codons upstream of the DNA binding domain (Fig. 1C). The human GR exists as multiple translation initiation variants, all of which are transcriptionally active, including the shortest, which lacks over 300 amino acids of N-terminal sequence contained in the full length GR^30^. One or more potential start codons remain available as in-frame initiation sites in all the previously reported exon 2 mutations of zebrafish *nr3c1* (Fig. 1), and thus those mutants produce mRNA that could potentially be translated into an N-terminally truncated GR. That such an isoform might retain activity as a transcription factor is shown by our observation that homozygous GR^369-^ zygotes injected with mRNA encoding a GR that lacks the first 310 amino acids of the full-length protein activate expression of the GR target *fkbp5*, albeit much less strongly than zygotes injected with mRNA encoding a full-length GR (Fig. 1E). In contrast, zygotes injected with mRNA encoding a GR lacking the first 369 amino acids and all start codons upstream of the DNA binding domain do not activate *fkbp5* (Fig. 1E). We conclude that GR^369-^ affords a complete loss of GR transcriptional function.

We used RNA-seq to determine how loss of GR function affects the larval transcriptome, both in larvae developed under normal conditions and in larvae treated chronically with cortisol. To overcome the low survival of embryos from homozygous mutant females and avoid nonspecific maternal effects that might be associated with poor egg quality, we took advantage of a visual background adaptation (VBA) screen to identify homozygous GR^369-^ progeny of a heterozygous cross, as those larvae lack a VBA response (VBA-larvae). Based on the observation that VBA-larvae comprised ∼¼ of the population, as expected for a recessive Mendelian trait, larvae that successfully mount a VBA response (VBA+ larvae) are predicted to consist of a 1:2 mixture of homozygous wild-type and heterozygous GR^369-^ mutants, and hence to contain at least one intact *nr3c1* allele encoding a functional GR. A principal component analysis of the RNA-seq data revealed that absence of a functional GR has a profound effect on gene expression in both normal and cortisol-treated larvae, such that transcriptomes from larvae lacking a GR are clearly distinguished from those that have one, and that cortisol treatment produces a coherent effect only in larvae with a functional GR (Fig. 2A). A pair-wise comparison of gene expression between VBA+ and VBA-larvae (accounting for PC2) identified with statistical significance about four hundred genes that are regulated by the GR in 5-day larvae at the time they were collected (midmorning). GO term enrichment analysis indicated that genes upregulated by the GR at that time are involved in metabolism and stress response, as would be expected, while genes downregulated by the GR are involved in epidermis development, cell adhesion, growth, and basement membrane formation, suggesting that the GR may function as a switch to downregulate those morphogenetic processes in late development, or to temporally segregate them to a certain time of day given the circadian dynamic of glucocorticoid signaling. Interestingly, numerous biological processes and individual genes affected by loss of GR function were similarly affected by chronic cortisol treatment in larvae with a GR. This suggests that one effect of the chronic treatment is to promote development of GR resistance, and moreover, that it does so via the GR. Such resistance might be construed as an adaptive response to the chronic exposure.

Comparing gene expression in VBA+ larvae treated with cortisol versus vehicle (accounting for PC1 in that experiment) identified over four thousand genes that are differentially expressed in response to chronic cortisol treatment. This latter number is substantially larger than the 555 differentially expressed genes identified in our previous analysis^23^ of the effects of chronic cortisol treatment and yielded a somewhat different result when subjected to GO term enrichment analysis (Figs. S5, S6). One major difference between the analyses reported here and that reported previously^23^ is that in the latter the embryos and larvae were cultured in the dark, whereas in the present study we cultured them from fertilization in a diurnal light-dark cycle. In zebrafish larvae the circadian clock is not synchronized until the fish are exposed to a light-dark cycle^42,43^, so our previous results may have had circadian asynchrony as a confounding variable. Indeed, the impact of this difference on the transcriptome is clearly seen in the PCA of the combined analysis of all three RNA-seq experiments (Fig. S14), accounting for nearly 20% of the variance. Despite this, the RNA-seq results reported here assessing effects of chronic cortisol exposure in wild-type and *klf9*^-/-^ larvae (Fig. 4) were from embryos developed with light-dark cycles, and in the wild type larvae produced an effect on proinflammatory gene expression similar to that of our earlier study, demonstrating that that effect was not an artifact of circadian asynchrony. Another difference between the both studies using wild-type larvae and that depicted in Figure 2 was that the latter measured transcriptomic effects of chronic cortisol treatment in a 1:2 mixture wild-type and heterozygous mutant larvae (VBA+); thus the results in VBA+ larvae would be expected to be less sensitive to any effects of the chronic exposure for which the GR is haplo-insufficient, a possibility supported by the comparison shown in Fig. 6B. Further work is required to more fully assess the effects of GR gene dosage on the transcriptome under both normal conditions and in response to chronic cortisol exposure.

The meta-analysis comparing all our RNA-seq experiments examining the transcriptomic effects of chronic cortisol treatment in wild-type or VBA+ larvae (Figs. 5 and 6 and Fig. S14) provides some important insights that are broadly relevant to RNA-seq data interpretation. One is that different experiments that examine the effects of a single variable under somewhat different conditions and in a limited number of biological replicates will often produce different lists of differentially expressed genes passing an arbitrary threshold of statistical significance (e.g. adjusted p <0.05). The reason is clear enough: biological systems are highly responsive to genetic and environmental factors that vary between experiments and affect gene expression, sometimes stochastically and/or in ways that are difficult to control. Nevertheless, GO analyses of the lists obtained from different experiments can detect consistent biological effects even if the gene lists differ in the individual genes that they include, particularly if methods such as GOrilla^44^ are used to test for statistically significant enrichment of GO terms toward one end or the other of a single list of genes ranked by some measurable criterion (e.g. a principal component that accounts for a specific condition as in Fig. 5). This underscores the important but often unappreciated point that statistical significance does not equate to biological significance, and generally is not a good sole criterion to assess the effects of a given condition on the expression of a given gene using high throughput methods such as RNA-seq. On the other hand, use of unbiased approaches such as PC analysis to parse the variance in the data can help identify robust condition-specific effects and provide insight into the biology underlying those effects when combined with GO term enrichment analysis. In the case of the experiments reported here this approach validated our earlier finding that chronic cortisol exposure leads to upregulation of pro-inflammatory gene expression and extended that result by showing that the upregulation depends on the GR target gene *klf9*.

Despite the large amount of variance in the RNA-seq results unrelated to the cortisol treatment, a small handful of genes identified as differentially expressed in our previous study were also identified as being differentially expressed in both of RNA-seq experiments presented in the present study, all being upregulated by the cortisol treatment (Fig. 6A). This set included *klf9*, one of only four genes showing GR-dependent expression in normal 5-day larvae that were also upregulated in VBA+ larvae in response to chronic cortisol, and only one of two that encode transcription factors, the other being *per1a* (Fig. 2B). Interestingly both *klf9* and *per1a* are involved in circadian regulation, and both have been shown to be GR targets in other vertebrate models^6-8,45^. Of these, only *klf9* was found to be upregulated in response to chronic cortisol treatment in all three of our RNA-seq experiments (Fig. 6A). Interestingly, *klf9* was also found to be the most commonly upregulated transcription factor in a recent meta-analysis of glucocorticoid-induced gene expression in the brain^46^. We have found by ATAC-seq that the promoter region of *klf9* is one of the most differentially open regions of chromatin in blood cells of adults derived from cortisol-treated embryos, possibly reflecting the chronically elevated cortisol levels maintained by those adults (Hartig et al., submitted). In mice *klf9* was recently shown to mediate glucocorticoid-induced metabolic dysregulation in liver^20^. Our RNA-seq results from *klf9*^-/-^ larvae (Fig. 4) show that Klf9 mediates the pro-inflammatory gene expression induced by chronic cortisol exposure that we reported previously^23^. Among other things Klf9 functions as a transcriptional repressor^10,11^, and in mouse macrophages as an incoherent feedforward regulator of the GR target *klf2*^12^, which functions to control inflammation^47^. Further work is needed to determine how *klf9* contributes to pro-inflammatory gene expression in response to chronic cortisol exposure, which could either be directly as a feedforward activator (possibly via effects on metabolism), indirectly as a feedforward repressor of an anti-inflammatory regulator like Klf2, or both. Our motif enrichment analysis of flanking sequences from the set of 149 genes upregulated by chronic cortisol in at least 2 of our 3 RNA-seq experiments indicated enrichment for KLF binding sites (Table S10); further work involving chromatin immunoprecipitation is needed to determine whether any of those sites are bound by Klf9 or Klf2.

Perhaps unsurprisingly, our results (Fig. 2 and Fig. S2) indicate that the GR is required for nearly all the transcriptomic effects of chronically elevated cortisol. However, eight genes upregulated by the treatment were found in the RNA-seq analysis to be upregulated in both VBA+ and VBA-larvae (Fig. 2B), indicating that the GR is not required for their upregulation. Interestingly most of these genes are known IEGs, and include the neuronal activity-dependent gene *npas4a*, the mammalian homologue of which is directly repressed by the GR^48^. One possible explanation is that the genes are upregulated by increased MR activity, which was recently shown to contribute to stress axis regulation in zebrafish larvae^28^. Further work in MR mutant fish^28^ will be needed to test this.

Finally, gene ontology analysis of genes upregulated by chronic cortisol treatment in VBA+ progeny of the GR^+/369-^ cross indicated a strong effect on biological processes associated with nervous system development and function. This is consistent with a recent report that injection of cortisol into eggs leads to increased neurogenesis in the larval brain^24^. In this regard it is interesting that *klf9* is a stress-responsive gene that regulates neural differentiation and plasticity^8,19^. The long-term dysregulation of the HPA/I axis caused by early life exposure to chronic stress and/or chronically elevated cortisol suggests that the exposure perturbs brain development and activity. Given its role in regulating plasticity in brain regions relevant to neuroendocrine function, it will be interesting to determine whether *klf9* contributes to those effects.

## Materials and Methods

### Zebrafish strains, husbandry, and embryo treatments

The AB wild-type strain was used for all genetic modifications. Husbandry and procedures were as described previously^23^. All animal procedures were approved by the Institutional Animal Care and Use Committee (IACUC) of the MDI Biological Laboratory, and all methods were performed in accordance with the relevant guidelines and regulations. Embryo culture and cortisol treatments were performed as previously described^23^, with one difference: embryos were cultured in a diurnal light-dark cycle (14 hours light – 10 hours dark). Briefly, fertilized eggs were collected in the morning, disinfected and at ∼4 hours post fertilization placed in dishes with either 1uM cortisol or vehicle (DMSO) added to embryo media. Embryos developed in a 28.5° C incubator with a 14/10 light/dark cycle synchronized with the core fish room. Media was changed daily.

### Construction of *nr3c1* and *klf9* mutant lines

To mutate *nr3c1* and *klf9* we used CRISPR-Cas9^49^, injecting zygotes with multiple guide RNAs for each gene and mRNA encoding Cas9. Guide RNAs were designed using the CHOP-CHOP algorithm^50,51^.

To generate the GR^369-^ mutant line fertilized wild-type AB embryos were injected at the 1 cell stage with 1-2 nL of a gRNA cocktail targeting *nr3c1* exons 2 and 3 (final concentration 40 ng/µL for each gRNA, 230 ng/µL Cas9 mRNA, 0.1 M KCl, and 0.01% phenol red indicator dye). Individual whole injected larvae were screened for mutations of the targeted regions by high resolution melt analysis (HRMA) of PCR amplicons containing those regions^52^. Detected mutations were then verified by Sanger sequencing of a PCR amplicon containing the targeted region. F0 adults bearing mutations were identified by HRMA of DNA extracted from tailfin clips, and germline mutations were then identified by PCR and HRMA/sequencing of sperm. F0 males with germline mutations were outcrossed to AB females, and heterozygous progeny were screened via sequencing from tailfin clips. The GR^369-^ mutation was identified and selected for by breeding over 2 additional generations to yield F3 homozygous progeny. Subsequent generations were maintained as heterozygotes for health and breeding purposes.

To generate the *klf9*^-/-^ mutant line fertilized wild-type AB embryos were injected at the 1-cell stage with <20% cell volume of injection mix consisting of 200ng/ul Cas9 mRNA, 100ng/ul guide RNA, 0.05% phenol red dye, and 0.2M KCl. Individual F0 larvae were screened with HRMA to confirm CRISPR efficacy. Larvae were placed into system and raised. Young adult fish were genotyped via HRMA using DNA extracted from fin clips, and mutations were confirmed by Sanger sequencing. F0 fish positive for mutation were outcrossed to wild-type (AB) fish. F1 offspring of this cross were screened by HRMA to confirm germline transmission, and Sanger sequencing identified the 2bp frame-shift mutation in one female founder. This female was out crossed with WT AB males. The resulting F2 fish were screened as young adults via fin clip and HRMA, and males positive for mutation were back crossed to the F1 founder female. Resulting F3 generation fish were screened via fin clip HRMA and sequenced to identify homozygous mutants as well as homozygous wild-type siblings. F3 generation was Mendelian 1:2:1 wild-type:heterozygote:homozygous-mutant ratio.

### Visual Background Adaptation (VBA) screen

To identify larvae lacking a functional GR a VBA screen was performed on 4-day old larvae as described^26^. Briefly, larvae were incubated for 20 minutes in a dark incubator then transferred to a white background and immediately examined under a stereomicroscope with brightfield optics. Larvae that failed to mount a VBA response were identified by the failure of melanophores to disperse, remaining clustered in a dark patch on the dorsal surface^26^. Larvae were segregated as VBA+ and VBA-cohorts and returned to culture for an additional day before they were collected for RNA-seq.

### RNA-seq and Data Analysis

At 3 hours zeitgeber time (post lights-on) on day 5 post-fertilization four biological replicates of cortisol-treated and control embryos were collected as follows. For the first RNA-seq experiment, one by one, a dish of larvae corresponding to a single condition was removed from the incubator, and 8 larvae per replicate (4 replicates) were collected in a 1.5 mL tube with minimal water and immediately snap frozen in liquid nitrogen. All replicates from one condition were collected sequentially before moving on to the next condition. Total RNA was extracted using the Qiagen RNA-Easy Plus mini kit (Qiagen). For the second RNA-seq experiment, four replicates of n=10 larvae were collected from a single dish for each condition and immediately snap frozen in liquid nitrogen. Collection of 16 samples occurred over 22 minutes. RNA was prepared as described above on two different days. On the first day (experimental replicates 1 and 2) the lysis buffer was added to all 8 frozen samples before homogenization, while on the second day the lysis buffer was added to each sample which was then immediately homogenized. This difference in sample preparation likely accounts for the large variance between samples prepared on day 1 and day 2. RNA was sent to the Oklahoma State Genomics Facility for Illumina library preparation and single-end sequencing.

RNA-seq libraries were generated with Illumina-compatible KAPA libraries and sequenced on an Illumina NextSeq 500 High Output sequencer. *klf9*^-/-^ and matched control samples were sequenced as single end 75-bp reads. VBA+ and VBA-samples were sequences as paired-end 75-bp samples.

Fastq formatted read files were preprocessed with Trimmomatic^53^ version 0.38 with default options, and then aligned to the Zebrafish genome version 11 as presented in Ensembl^54^ version 93, using the STAR aligner^55^ version 2.6.1b. The Ensembl transcriptome was preprocessed with a splice junction overhang of 100 nt. Following alignment, the resulting BAM files were processed with RSEM^56^ version 1.3.0 for isoform and gene-level expression estimates. The resulting gene-level expression values were merged into a single expression matrix with an in-house python script. EDASeq^57^ carried out in R version 3.6.1 was used to further normalize data for systematic effects, using gene-level length and GC-content as downloaded from Ensembl version 98 using EDAseq’s included scripts. “WithinLane” normalization with GC content was judged as superior to that based on length. Final normalized gene-level counts (which = “full”) were generated using GC-based WithinLaneNormalization followed by BetweenLaneNormalization. Subsequent differential expression analysis was carried out in R version 3.6.1 with the DESeq2^58^ version 1.24.0. DESeq2 was also used to generate a rlog-matrix which was subsequently Z-transformed to normalize each gene across all samples. The Z-transformed rlog matrix was then loaded into JMP version 15 and used for PCA (using the “Wide Method”) after thresholding to an average rlog expression value of 7.5.

Both RNA-seq dataset have been deposited in the NCBI Gene Expression Omnibus database, under accession numbers GSE144884 (GR+/- experiment) and GSE144885 (Klf9+/- experiment).

The previously published RNA-seq data set was reprocessed as just described, and an overall matrix of only wild-type or VBA+ samples was generated and jointly normalized with EDASeq and then subjected to PCA analysis as described in the previous paragraph.

DESeq2 output tables were filtered with the following restrictions for the identification of statistically significant differentially expressed genes: BaseMean >= 75 (average of 5 counts per sample), padj <= 0.05, |log2foldchange| >= 0.5, estimated standard error of the log2foldchange < 1.0.

The GOrilla algorithm^44^ (http://cbl-gorilla.cs.technion.ac.il/) was used for Gene Ontology term enrichment analysis, and the data were visualized using REVIGO^59^ (http://revigo.irb.hr/). Venn diagrams were generated using Venny 2.1 (https://bioinfogp.cnb.csic.es/tools/venny/). HOMER motif enrichment analysis^37^ was used to compare incidence of known vertebrate motifs in a list of promoters of interest with incidence in a background list of all zebrafish promoters by running the findMotifs program (http://homer.ucsd.edu/homer/microarray/index.html) using default settings except that sequence from - 1500bp to +500 bp relative to the transcription start site was searched for motifs from 10bp to 18bp in length.

### Quantitative Reverse Transcription and Polymerase Chain Reaction (qRT-PCR)

Total RNA purified from snap-frozen larvae using the Trizol method and used as template to synthesize random-primed cDNA using the Primescript cDNA synthesis kit (TaKaRa). Relative gene expression levels were measured by qRT-PCR, using the delta-delta Ct method as described previously^23^, and *eif5a* as a reference gene. Examination of the results of multiple RNA-seq data sets indicated that *eif5a* activity was highly stable across treatments and genotypes. In many experiments beta-actin was also used as a reference gene, and this did not substantially change the results.

## Acknowledgements

This work was supported by grants from the National Institutes of Health (R03-HD099468, P20-GM104318, and P20-GM103423), by a Morris Scientific Discovery Award to JAC, and by a gift from The Linde Packman Lab for Biosciences Innovation to Colby College.

## Author Contributions

IG and EIH performed most of the experiments and did the matings, microinjections and husbandry to establish the *klf9* and *nr3c1* mutant zebrafish strains; SZ designed the gRNAs and performed the initial PCR and sequencing analyses to identify fish containing the desired *nr3c1* mutations; IG performed the mRNA injection experiments, qRT-PCR, RNA-seq data analysis, and HOMER motif enrichment analysis; ART contributed to the RNA-seq experimental design and data analysis; LNH wrote the code used in the RNA-seq data analysis; NJM performed the initial analysis of the RNA-seq data; JHG directed the computational analyses and performed the PC and DGE analyses; JAC conceived the study, contributed to analysis of the data and drafted the manuscript.

## Notes

#### Summary of Updates

This revision has a revised Figure 4, new Figures 5 and 6, additional supplementary material, and some new text in the Discussion.

