## Supplemental Figures for "Klf9 is a key feedforward regulator of the transcriptomic response to glucocorticoid receptor activity"

```
Human              1 MDSKESLTPGREENPSSVLAQERGDVMDFYKTLRGGATVKVSASSPSLAV     50
                           :..|..||...  ..||.:.:|:.|....|...:...|:.|:|.
Zfish              1 ------MDQGGLENGKK--RDERLNTLDYNKRATEGILPRRIQSTMSVAP     42

Human             51 AS---QSDSKQRRLLVDFPKGSVSNAQQPDLSKAVSLSMGLYMGETETKV     97
                     .|   |:....:.:..|.|.| :||:  |.|.:..|....:..::.|.|:
Zfish             43 TSMVPQAGPMMQPVSGDIPNG-LSNS--PTLEEHTSSVSSIFGDDSELKL     89

Human             98 MGND-LGFPQQGQISLSSGETDLKLLEESIANLNRSTSVPENPKSSASTA    146
                     :|.: ....||..:..:.|:: |..||.|||:||       ||..|..:.
Zfish             90 LGKEQRALQQQTLVPFTLGDS-LSGLEASIADLN-------NPSPSMDSL    131

Human            147 VSAA-------PTEKEFPKTHSDVSSEQQHLKGQTG--TNGGNVKLYTTD    187
                     :...       .||...|....|:..:|... |..|  .:.||.||::
Zfish            132 IGGVDPNLFPLKTEDFSPMIKGDMDLDQDSF-GHIGKDVDVGNHKLFS--    178

Human            188 QSTFDILQDLEFSSGSPGKETNESPWRSDLLIDENCLLSPLAGEDDSFLL    237
                     .:|.|:|||.|. .|||          ||..:.::..||.:.  :|:.|.
Zfish            179 DNTLDLLQDFEL-DGSP----------SDFYVADDAFLSTIG--EDALLS    215

Human            238 EGNSNEDCKPLILPDTKPKIKDNGDL------VLSSPSNVTLPQVKTEKE    281
                     |..:|.|      .|:|..:..:..|      .||:.:...||.:|.||:
Zfish            216 ELPTNLD------RDSKAAVSGSNTLNGTASSSLSTANTSILPNIKVEKD    259

Human            282 DFIELCTPGVIKQEKLGTVYCQASFPGANIIGNKMSAISVHGVSTSGGQM    331
                     ..|:||||||||||..|..|||.        |...:.|::.||:||.||.
Zfish            260 SIIQLCTPGVIKQENTGASYCQG--------GLHSTPINICGVTTSSGQS    301

Human            332 YHYDMN--TASLSQQQDQKPIFNVIPPIPVGSENWNRCQGSGD-DNLTSL    378
                     :.:..:  ||.:..|:||||.||:..|:....:.|:|.||.|: ..:...
Zfish            302 FLFGNSSPTAVVGLQKDQKPDFNMYTPLTSSGDGWSRSQGFGNVSGMQQR    351

Human            379 GTLNFPGRTVFSNGYSSPSMRPDVSSPPSSSSTATTGPPPKLCLVCSDEA    428
                     .:|.|.     .|..|||..||: .|..:||:...|| ..|:||||||||
Zfish            352 ASLCFS-----KNFSSSPYSRPE-DSTATSSAGGKTG-THKICLVCSDEA    394

Human            429 SGCHYGVLTCGSCKVFFKRAVEGQHNYLCAGRNDCIIDKIRRKNCPACRY    478
                     |||||||||||||||||||||||||||||||||||||||||||||||||:
Zfish            395 SGCHYGVLTCGSCKVFFKRAVEGQHNYLCAGRNDCIIDKIRRKNCPACRF    444

Human            479 RKCLQAGMNLEARKTKKKIKGIQQATTGVSQETSEN------PGNKTIVP    522
                     ||||.|||||||||:|.|.:   ||...:.|::...      |..:.:||
Zfish            445 RKCLMAGMNLEARKSKSKAR---QAGKVIQQQSIPERNLPPLPEARALVP    491

Human            523 ATLPQLTPTLVSLLEVIEPEVLYAGYDSSVPDSTWRIMTTLNMLGGRQVI    572
                     ..:|||.||::|||:.|||:.|||||||::||::.|:|||||.|||||||
Zfish            492 KPMPQLVPTMLSLLKAIEPDTLYAGYDSTIPDTSVRLMTTLNRLGGRQVI    541

Human            573 AAVKWAKAIPGFRNLHLDDQMTLLQYSWMFLMAFALGWRSYRQSSANLLC    622
                     :|||||||:||||||||||||||||.||:|:|:|.||||||:..:.|:||
Zfish            542 SAVKWAKALPGFRNLHLDDQMTLLQCSWLFIMSFGLGWRSYQHCNGNMLC    591

Human            623 FAPDLIINEQRMTLPCMYDQCKHMLYVSSELHRLQVSYEEYLCMKTLLLL    672
                     |||||:|||:||.||.|.|||:.||.:|:|..|||||.|||||||.||||
Zfish            592 FAPDLVINEERMKLPYMSDQCEQMLKISNEFVRLQVSTEEYLCMKVLLLL    641

Human            673 SSVPKDGLKSQELFDEIRMTYIKELGKAIVKREGNSSQNWQRFYQLTKLL    722
                     ::|||||||||.:|||:||:|||||||||||||.||||||||||||||||
Zfish            642 NTVPKDGLKSQSVFDELRMSYIKELGKAIVKREENSSQNWQRFYQLTKLL    691

Human            723 DSMHEVVENLLNYCFQTFLDKTMSIEFPEMLAEIITNQIPKYSNGNIKKL    772
                     ||||::|..|||:||.||::|::|:||||||||||:||:||:.:|::|.|
Zfish            692 DSMHDLVGGLLNFCFYTFVNKSLSVEFPEMLAEIISNQLPKFKDGSVKPL    741

Human            773 LFHQK    777
                     |||||
Zfish            742 LFHQK    746
```

**Figure S1. Needle amino acid sequence alignment of human and zebrafish glucocorticoid receptors.** Methionines available as potential N-termini via alternative translation initiation are highlighted. Those in the zebrafish sequence were identified by the NetStart 1.0 algorithm. Those in the human GR are known to produce transcriptionally functional products (Lu and Cidlowski, 2005). The DNA binding domain is underlined.

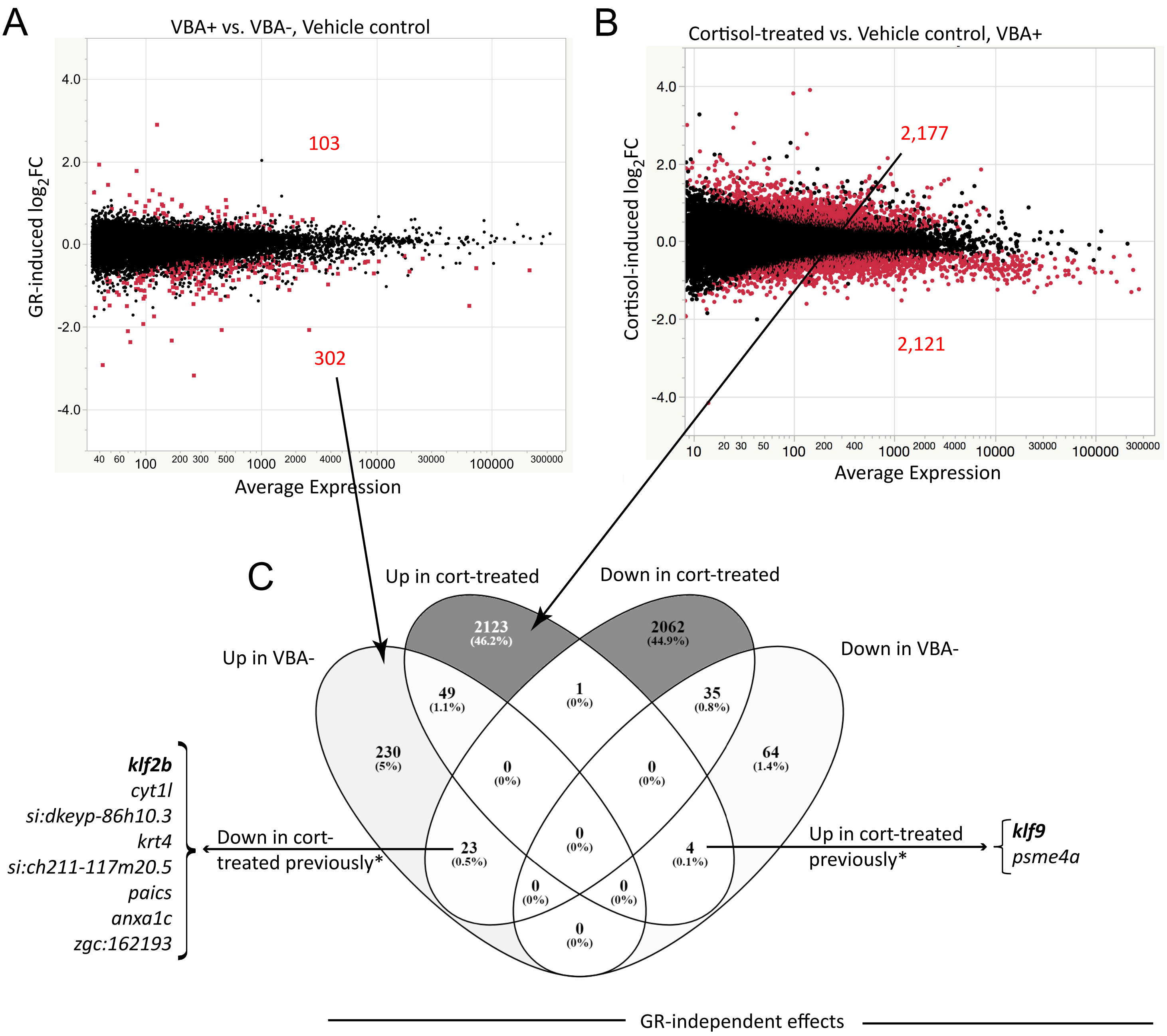


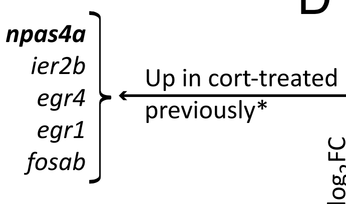


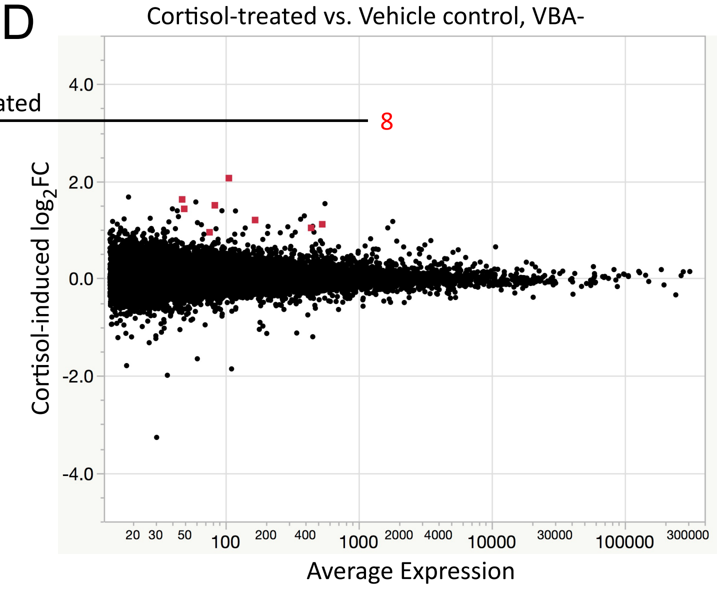


**Figure S2. Transcriptomic effects of GR loss of function and chronic cortisol exposure in larvae containing or lacking a GR.** (A) MA plot comparing gene expression in VBA+ and VBA- larvae developed under control conditions; (B) MA plot comparing gene expression in cortisol-treated VBA+ and vehicle-treated VBA+ larvae; (C) Venn diagram comparing the results of (A) and (B), and showing lists of genes* previously found to be down- or up-regulated by chronic cortisol (*Hartig et al., 2016); (D) MA plot comparing gene expression in cortisol-treated VBA- and vehicle-treated VBA- larvae. Diagram generated by Venny (https://bioinfogp.cnb.csic.es/tools/venny/).

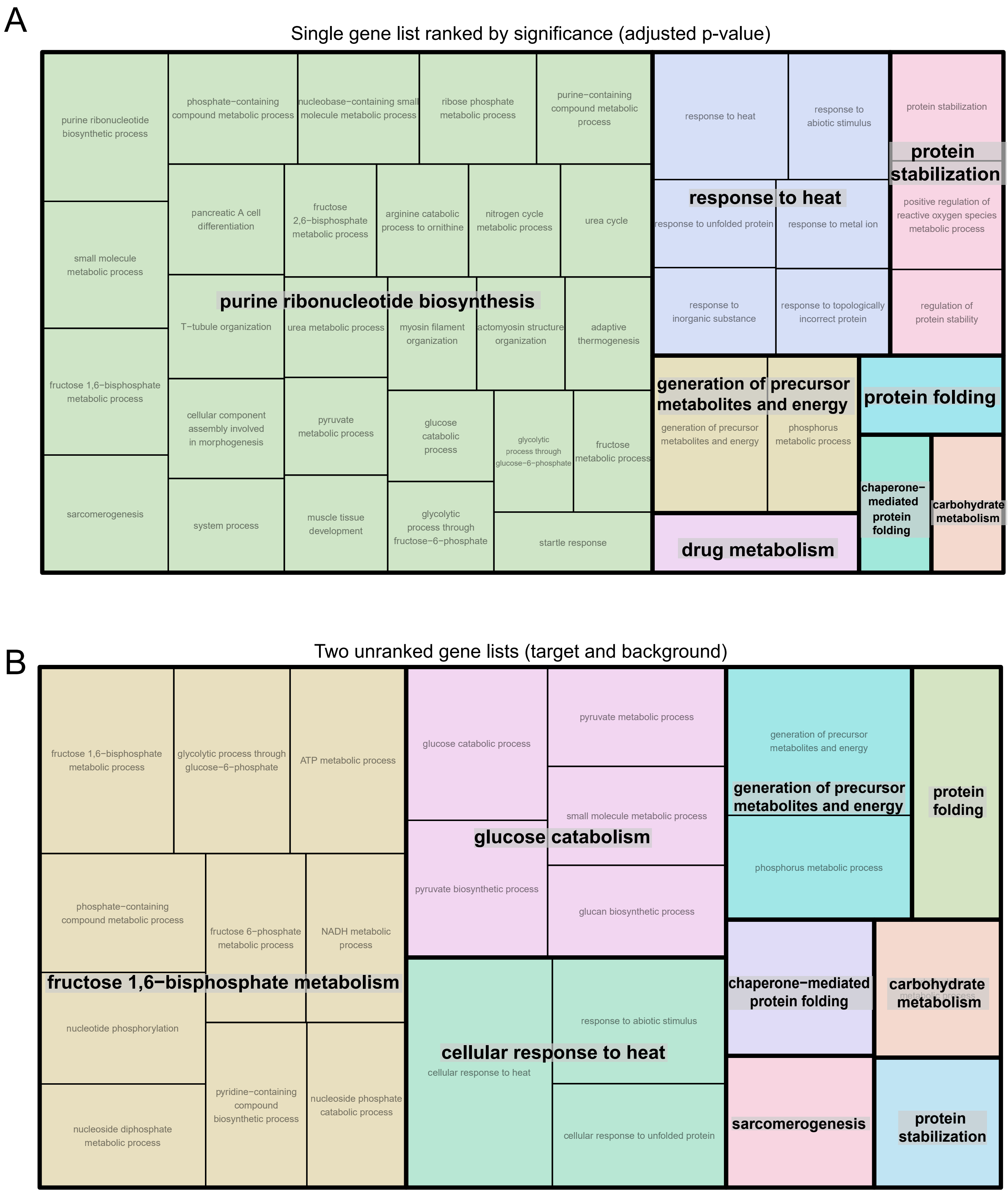


**Figure S3. REVIGO Gene Ontology treemaps of processes upregulated by the GR under normal conditions**, identified by GOrilla analysis of (A) single ranked gene list and (B) two unranked gene lists (target and background, with target genes having an adjusted p-value <0.05) obtained from differential gene expression analysis.

A

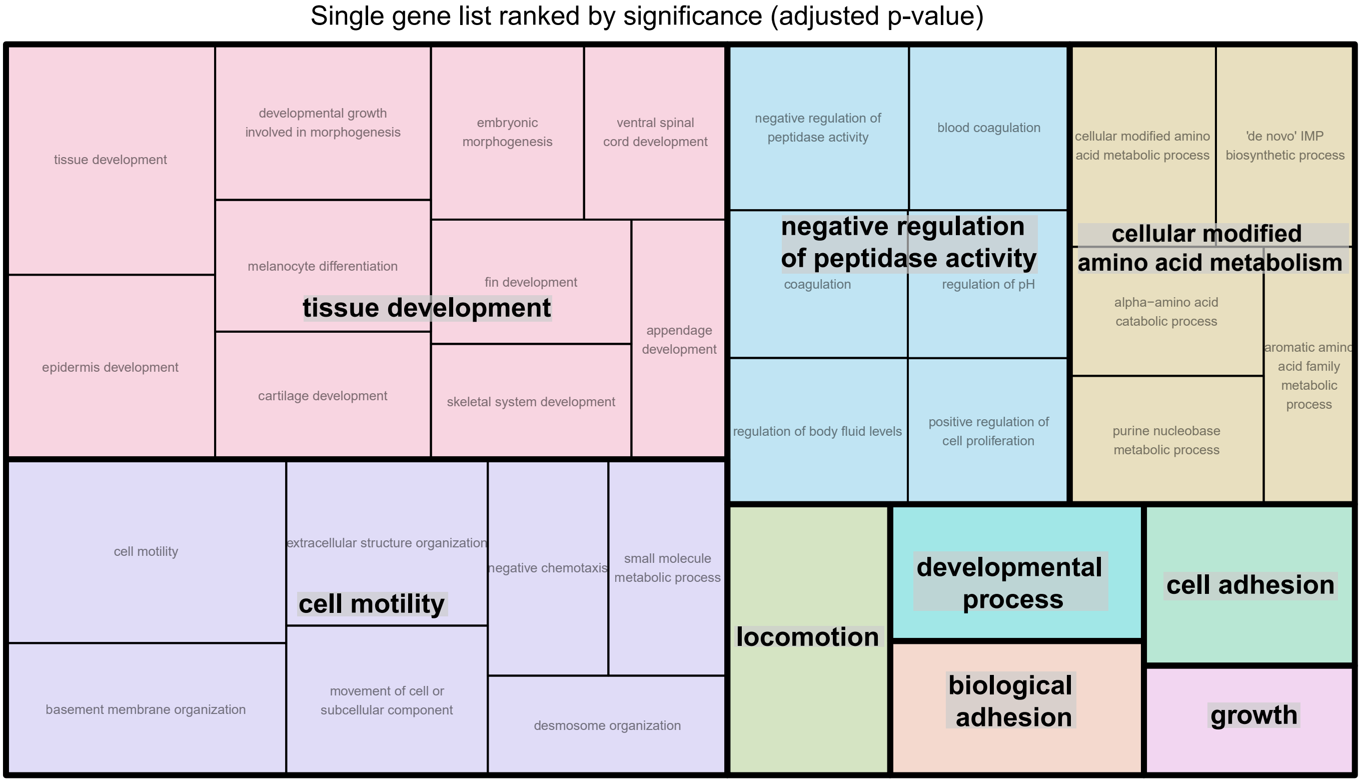


B

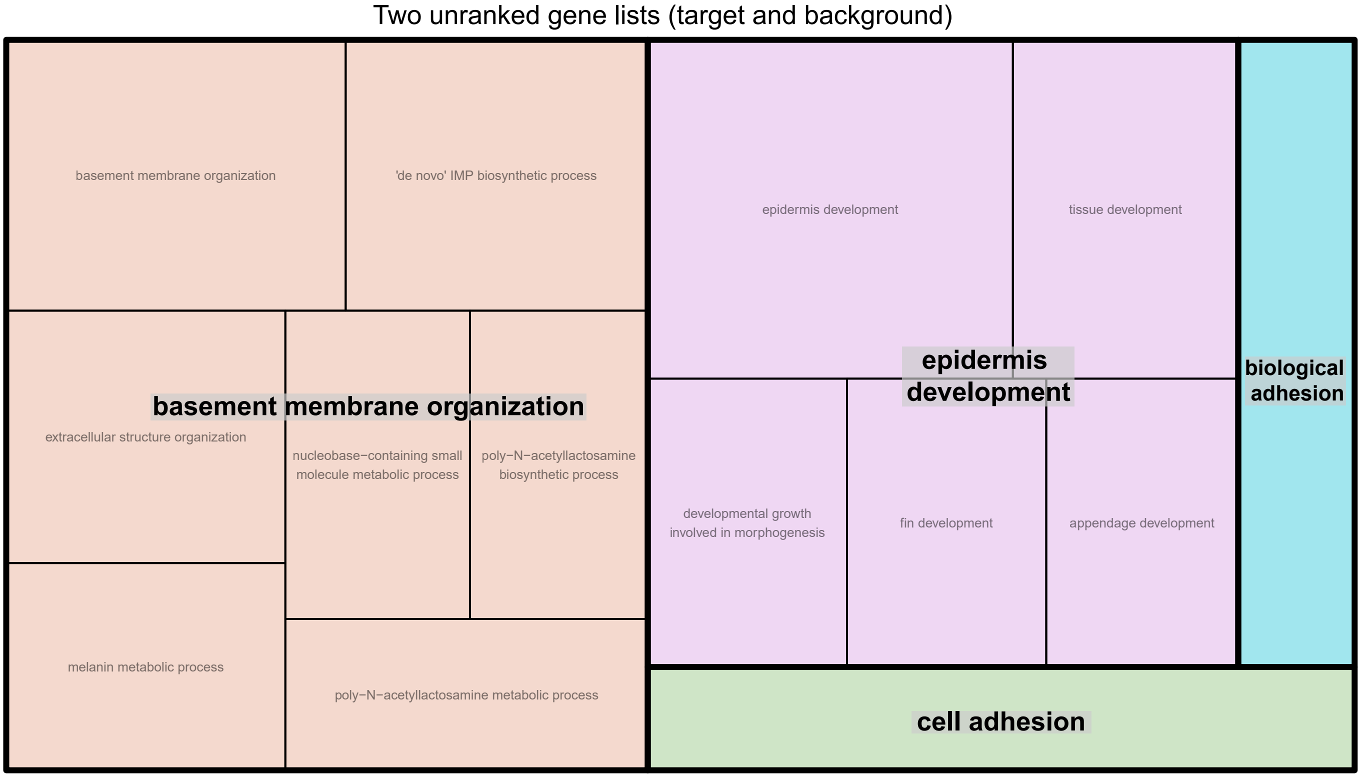


**Figure S4. REVIGO Gene Ontology treemaps of processes downregulated by the GR under normal conditions**, identified by GOrilla analysis of (A) single ranked gene list and (B) two unranked gene lists (target and background, with target genes having an adjusted p-value <0.05) obtained from differential gene expression analysis.

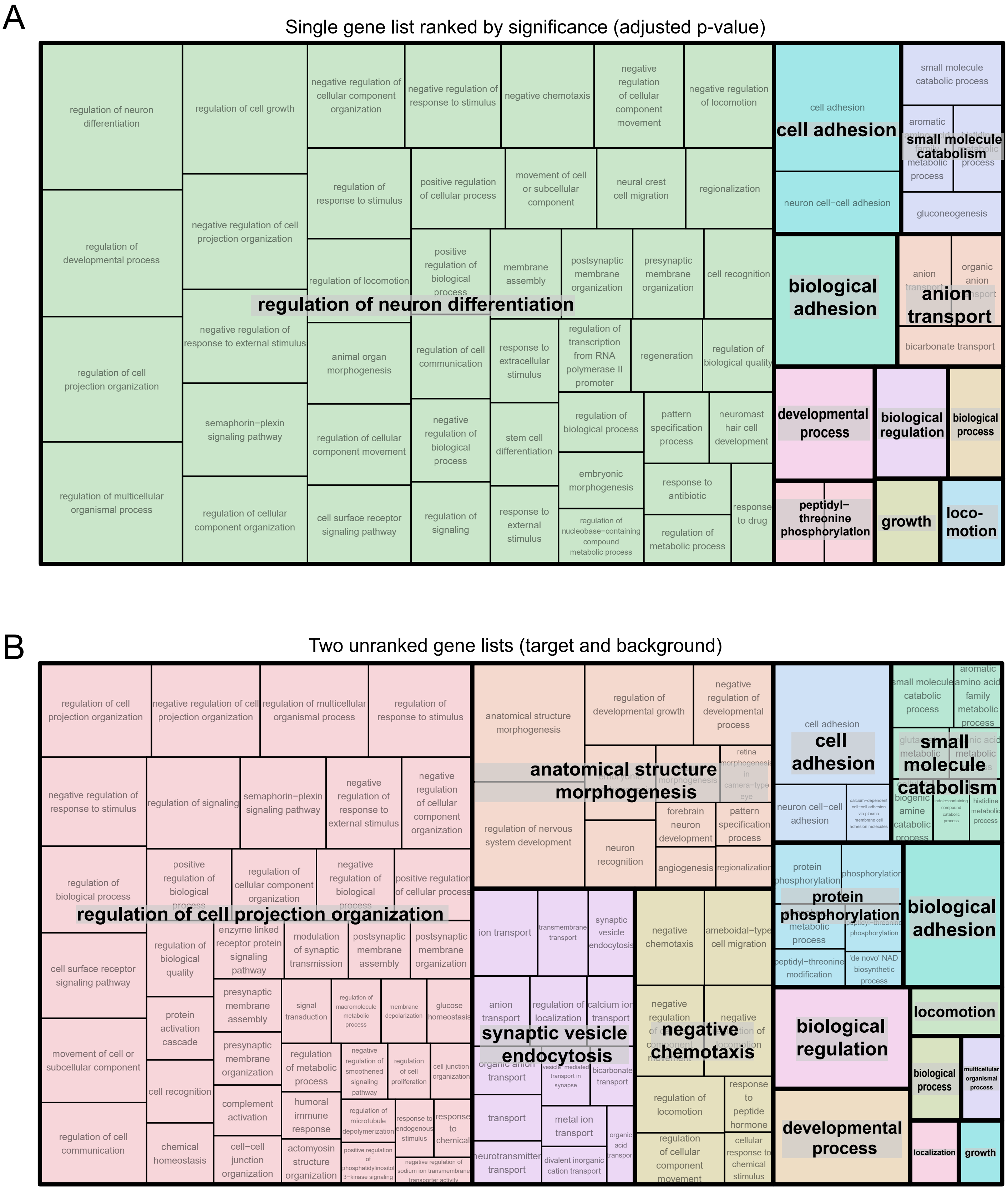


**Figure S5. REVIGO Gene Ontology treemaps of processes upregulated by chronic cortisol treatment in larvae with a GR,** identified by GOrilla analysis of (A) single ranked gene list and (B) two unranked gene lists (target and background, with target genes having an adjusted p-value <0.05) obtained from differential gene expression analysis.

A

Single gene list ranked by significance (adjusted p-value)

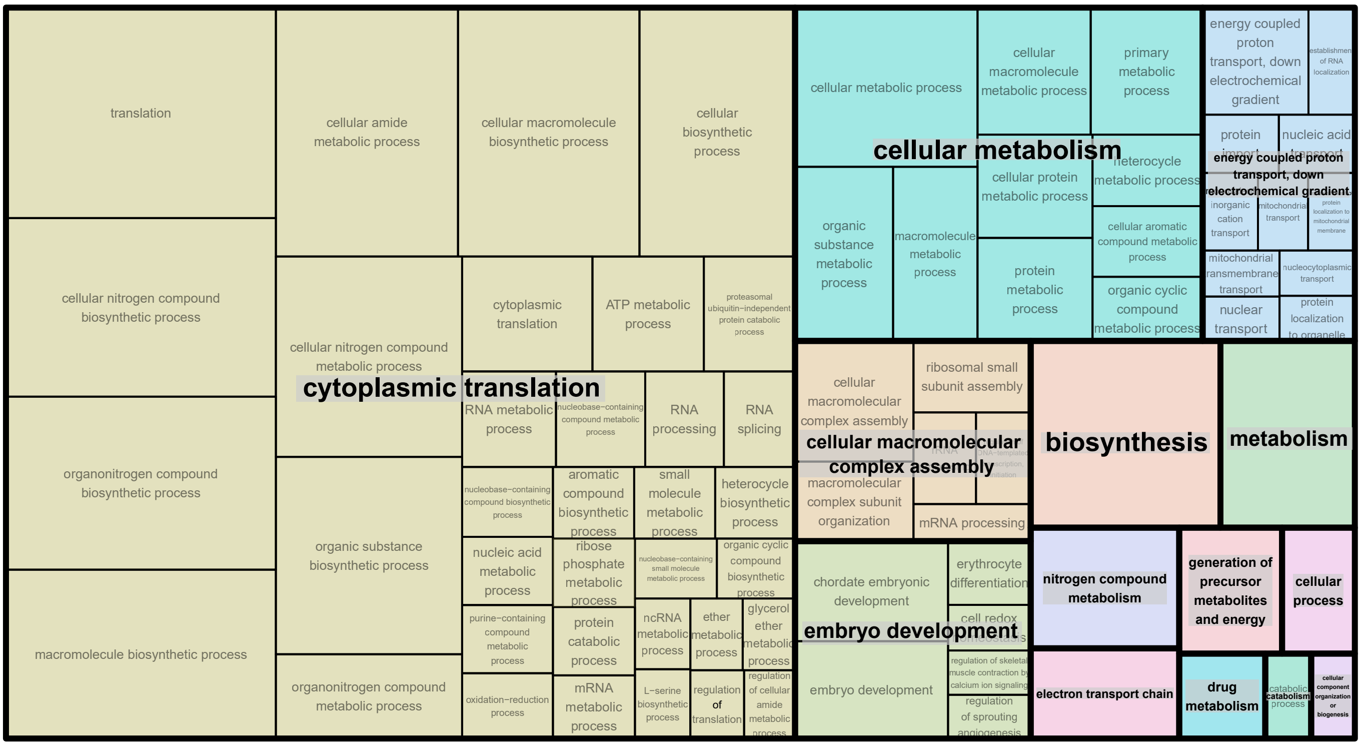


B

Two unranked gene lists (target and background)

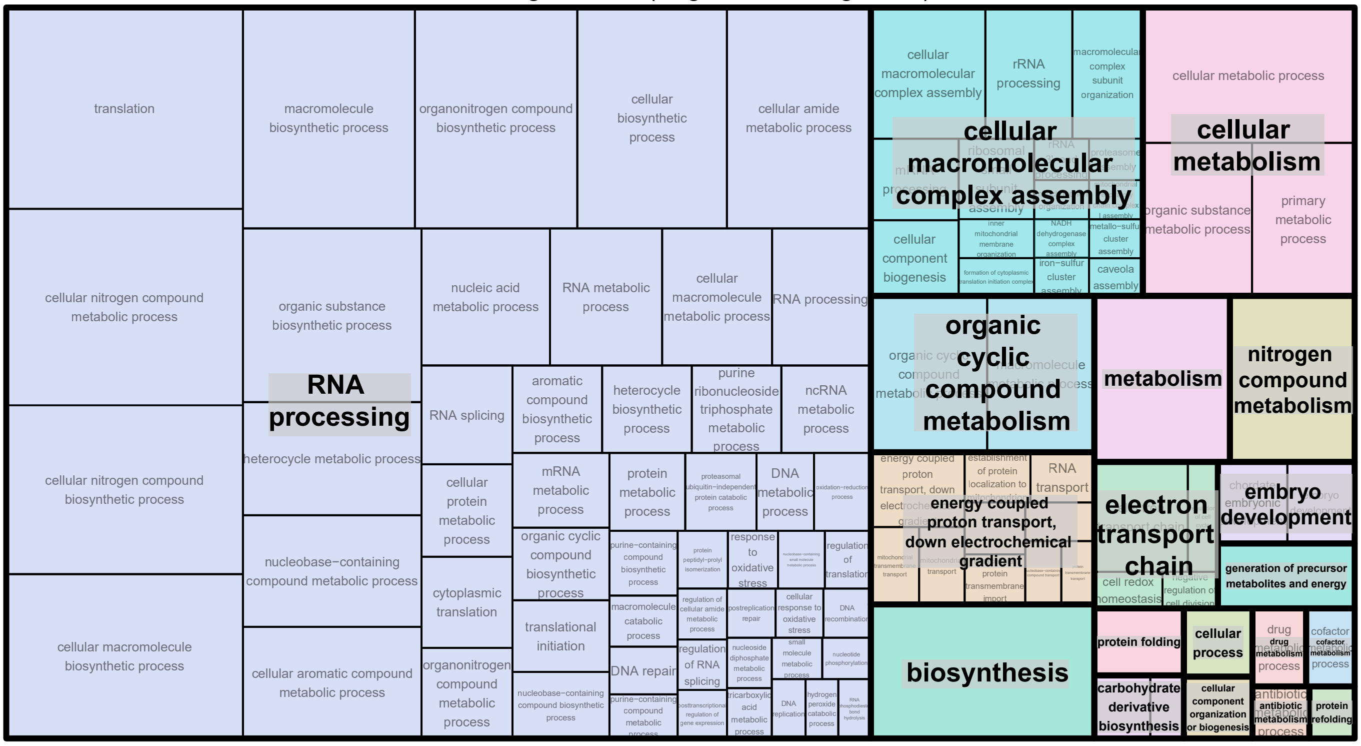


**Figure S6. REVIGO Gene Ontology treemaps of processes downregulated by chronic cortisol treatment in larvae with a GR,** identified by GOrilla analysis of (A) single ranked gene list and (B) two unranked gene lists (target and background, with target genes having an adjusted p-value <0.05) obtained from differential gene expression analysis.

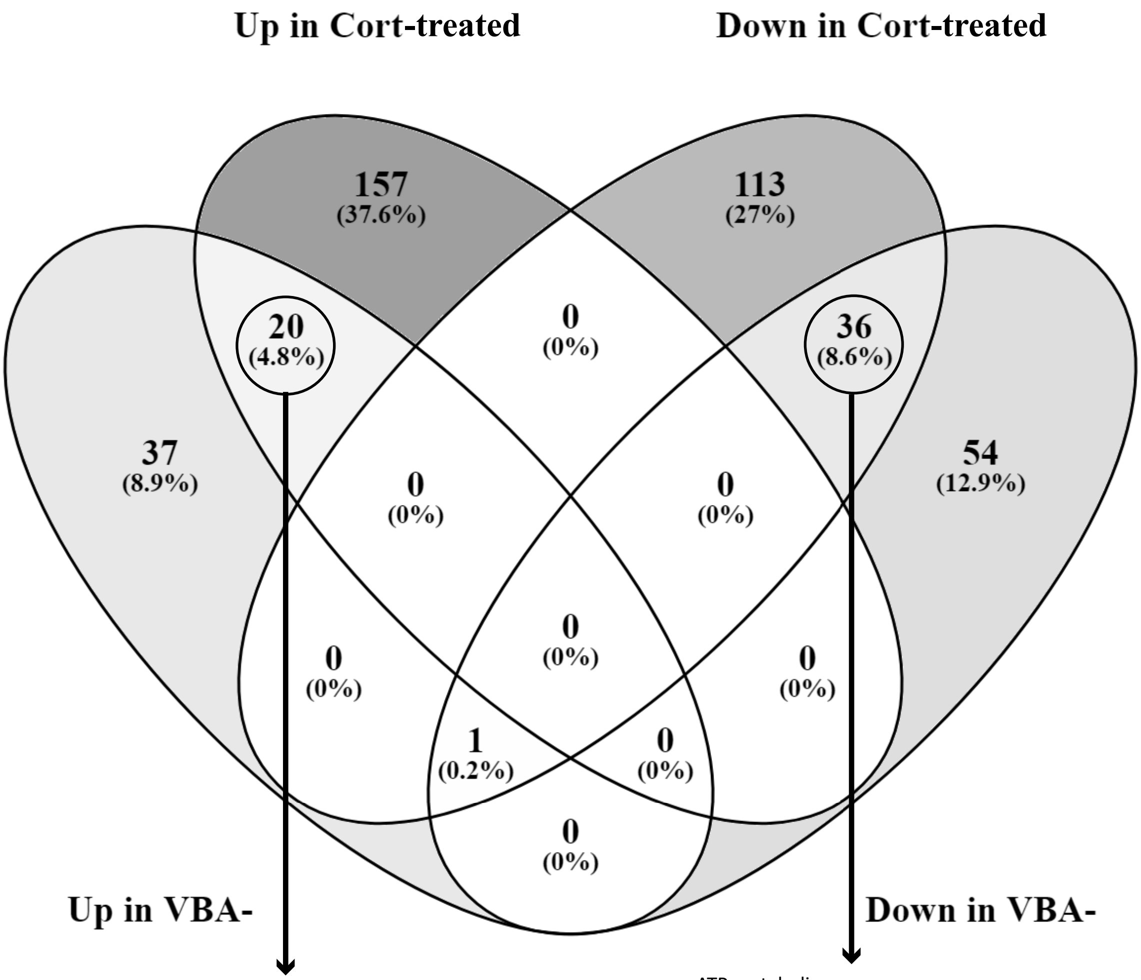


biological adhesion
cell adhesion
anatomical structure morphogenesis
developmental process
negative regulation of growth
negative chemotaxis
anatomical structure development
movement of cell or subcellular component
ameboidal-type cell migration
growth
developmental growth
cell migration
locomotion
cell motility
tissue development
cellular amino acid catabolic process
embryonic morphogenesis
alpha-amino acid catabolic process
aromatic amino acid family metabolic process
cell-substrate adhesion

ATP metabolic process
purine ribonucleoside triphosphate metabolic process
purine nucleoside triphosphate metabolic process
ribonucleoside triphosphate metabolic process
ATP biosynthetic process
purine ribonucleoside triphosphate biosynthetic process
purine nucleoside triphosphate biosynthetic process
nucleoside triphosphate metabolic process
purine nucleoside monophosphate biosynthetic process
purine ribonucleoside monophosphate biosynthetic process
purine nucleoside monophosphate metabolic process
purine ribonucleoside monophosphate metabolic process
nucleoside triphosphate biosynthetic process
ribonucleoside triphosphate biosynthetic process
ribonucleoside monophosphate biosynthetic process
nucleoside monophosphate biosynthetic process
ribonucleoside monophosphate metabolic process
purine ribonucleotide biosynthetic process
nucleoside monophosphate metabolic process
purine-containing compound biosynthetic process
purine nucleotide biosynthetic process
ribonucleotide biosynthetic process
drug metabolic process
ribose phosphate biosynthetic process
purine ribonucleotide metabolic process
nucleotide biosynthetic process
ribonucleotide metabolic process
purine-containing compound metabolic process
nucleoside phosphate biosynthetic process
purine nucleotide metabolic process
ribose phosphate metabolic process
nucleotide metabolic process
nucleoside phosphate metabolic process
nucleobase-containing small molecule metabolic process
organophosphate biosynthetic process

**Figure S7. Venn diagram of numbers of biological processes shown by GOrilla analysis to be affected by presence or absence of the GR and by chronic cortisol treatment and overlap between them.** Diagram generated by Venny (https://bioinfogp.cnb.csic.es/tools/venny/).

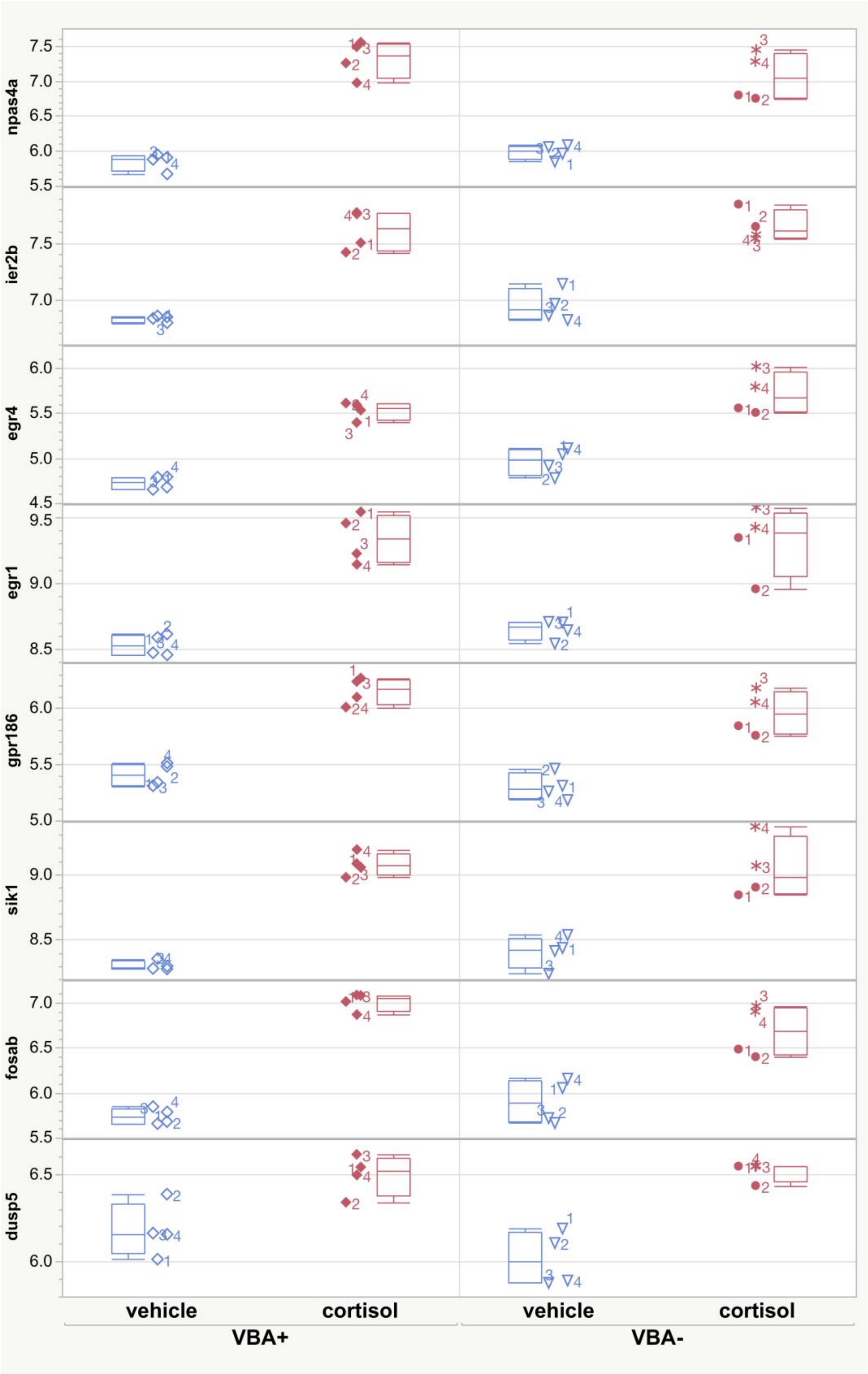


**Figure S8. Expression levels of eight genes that are upregulated by chronic cortisol treatment in both VBA+ and VBA- larvae, measured by RNA-seq.** Several of these genes (*npas4a, fosab, egr1, egr4, ier2b*) were found in our previously reported RNA-seq analysis to be upregulated in wild-type larvae treated chronically with cortisol (Hartig et al., 2016).

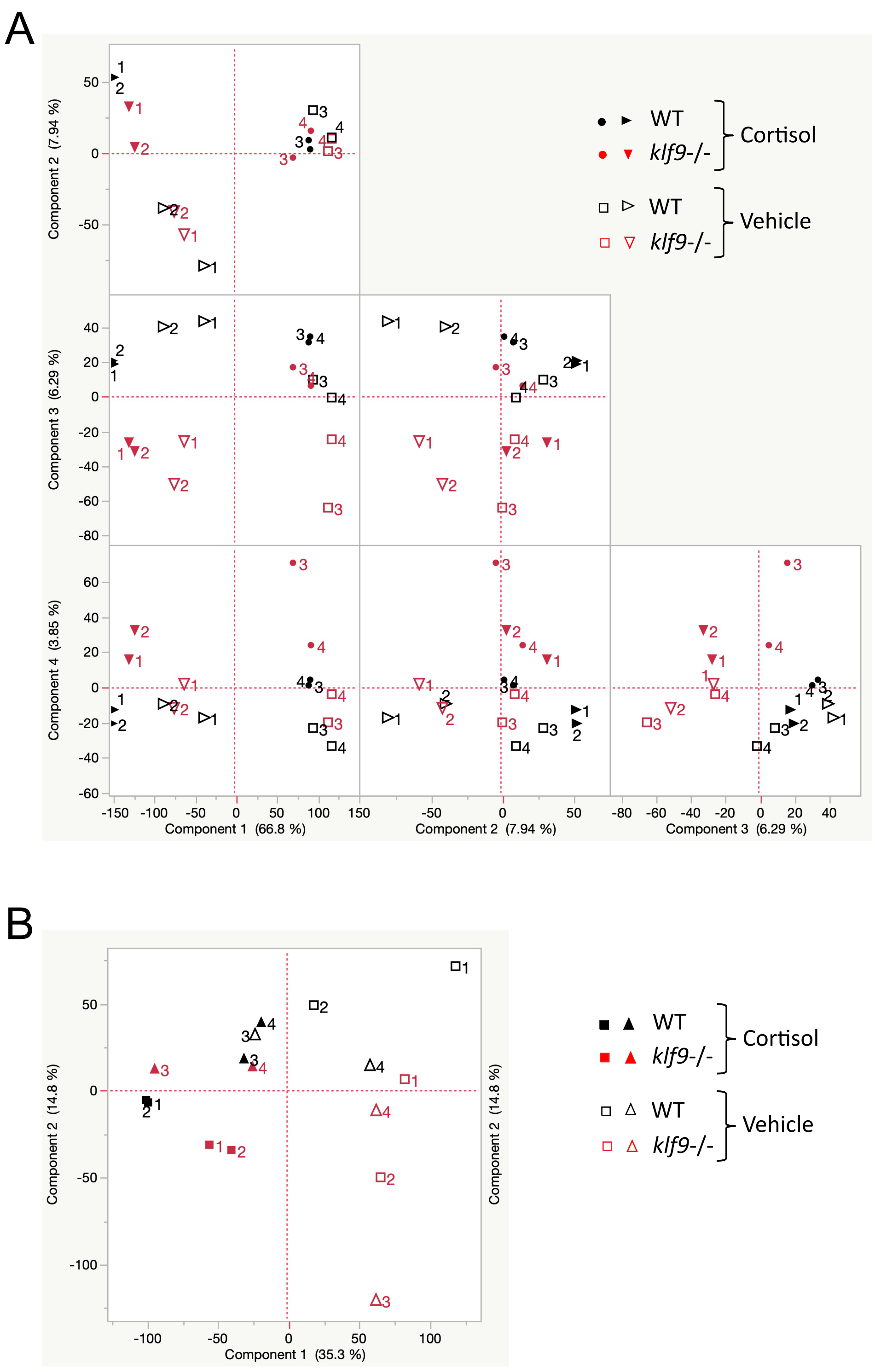


**Figure S9. Principal component (PC) plots of the RNA-seq data comparing transcriptomes of 5 day old wild-type (WT) and *klf9*$^{-/-}$ larvae developed normally (vehicle) or with chronic cortisol treatment.** (A) Plots of the first four PCs when all samples were rlog normalized as a group. (B) Plots of the first two PCs obtained when the data were rlog normalized independently for each day the samples were prepared (day 1, replicates 1 and 2; day 2, replicates 3 and 4).

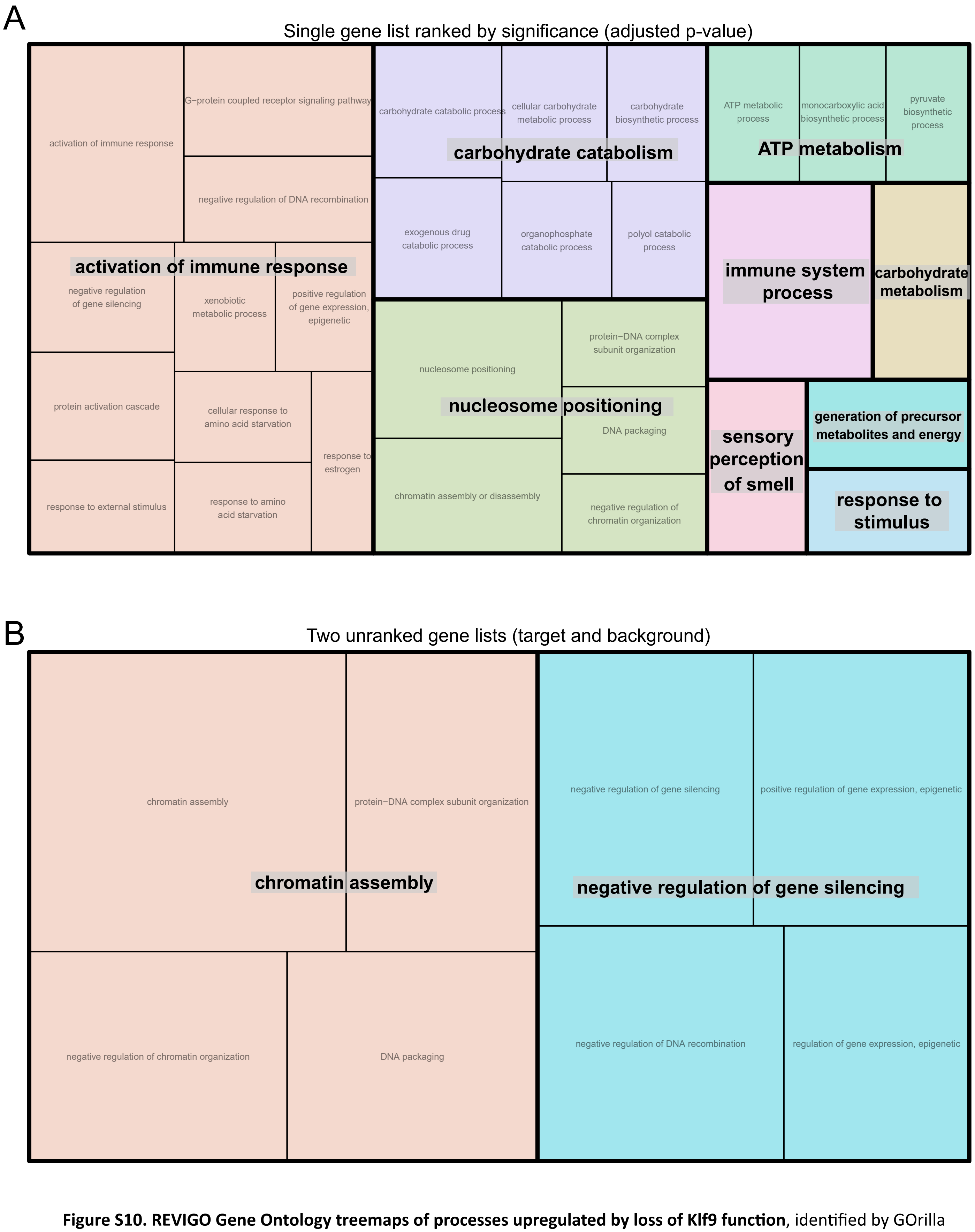


**Figure S10. REVIGO Gene Ontology treemaps of processes upregulated by loss of Klf9 function**, identified by GOrilla analysis of (A) single ranked gene list and (B) two unranked gene lists (target and background, with target genes having an adjusted p-value <0.05) obtained from differential gene expression analysis.

A

Single gene list ranked by significance (adjusted p-value)

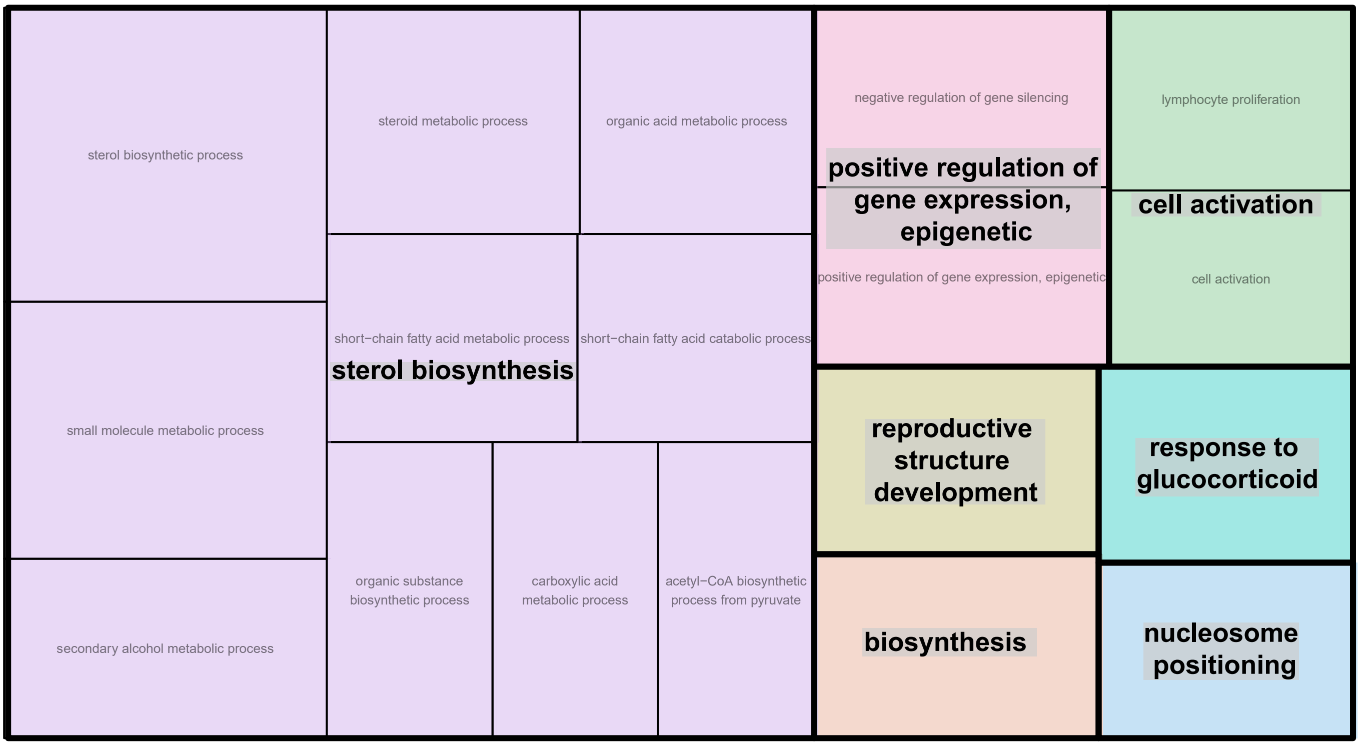


B

Two unranked gene lists (target and background)

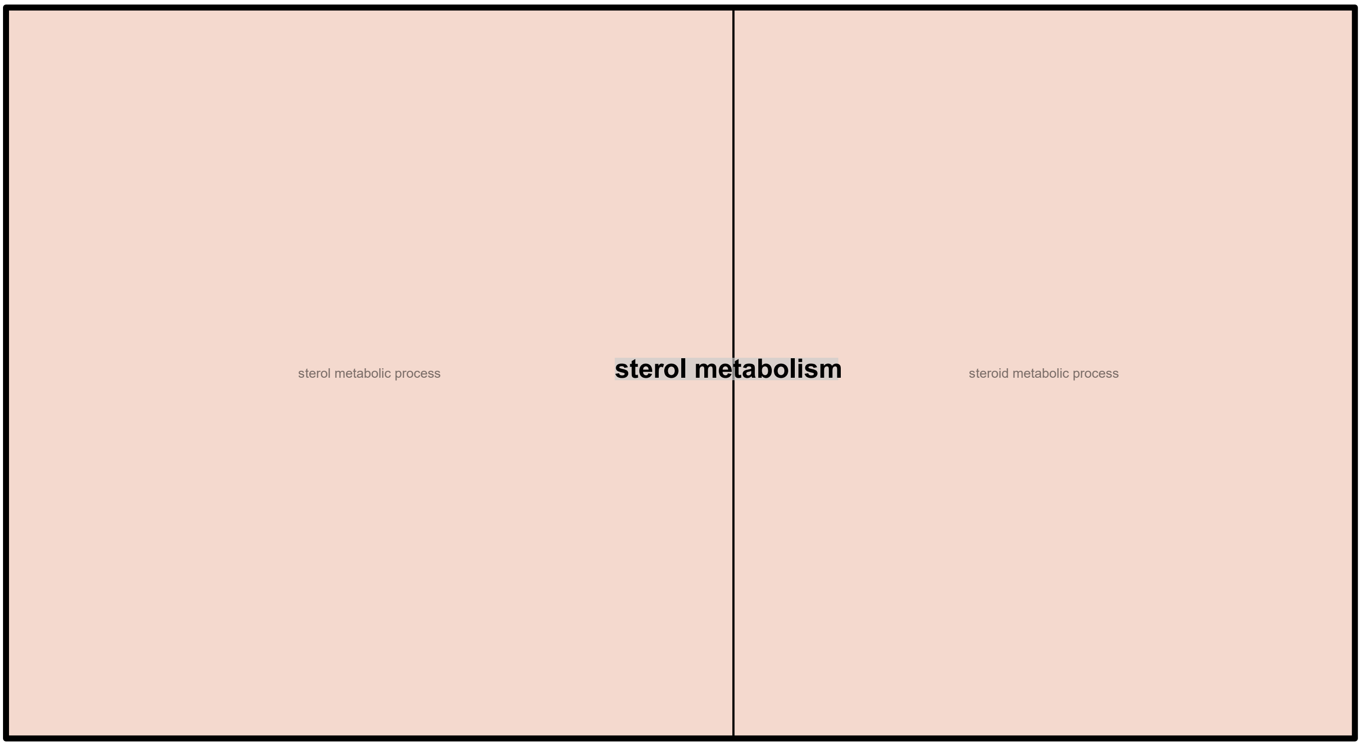


**Figure S11. REVIGO Gene Ontology treemaps of processes downegulated by loss of Klf9 function**, identified by GOrilla analysis of (A) single ranked gene list and (B) two unranked gene lists (target and background, with target genes having an adjusted p-value <0.05) obtained from differential gene expression analysis.

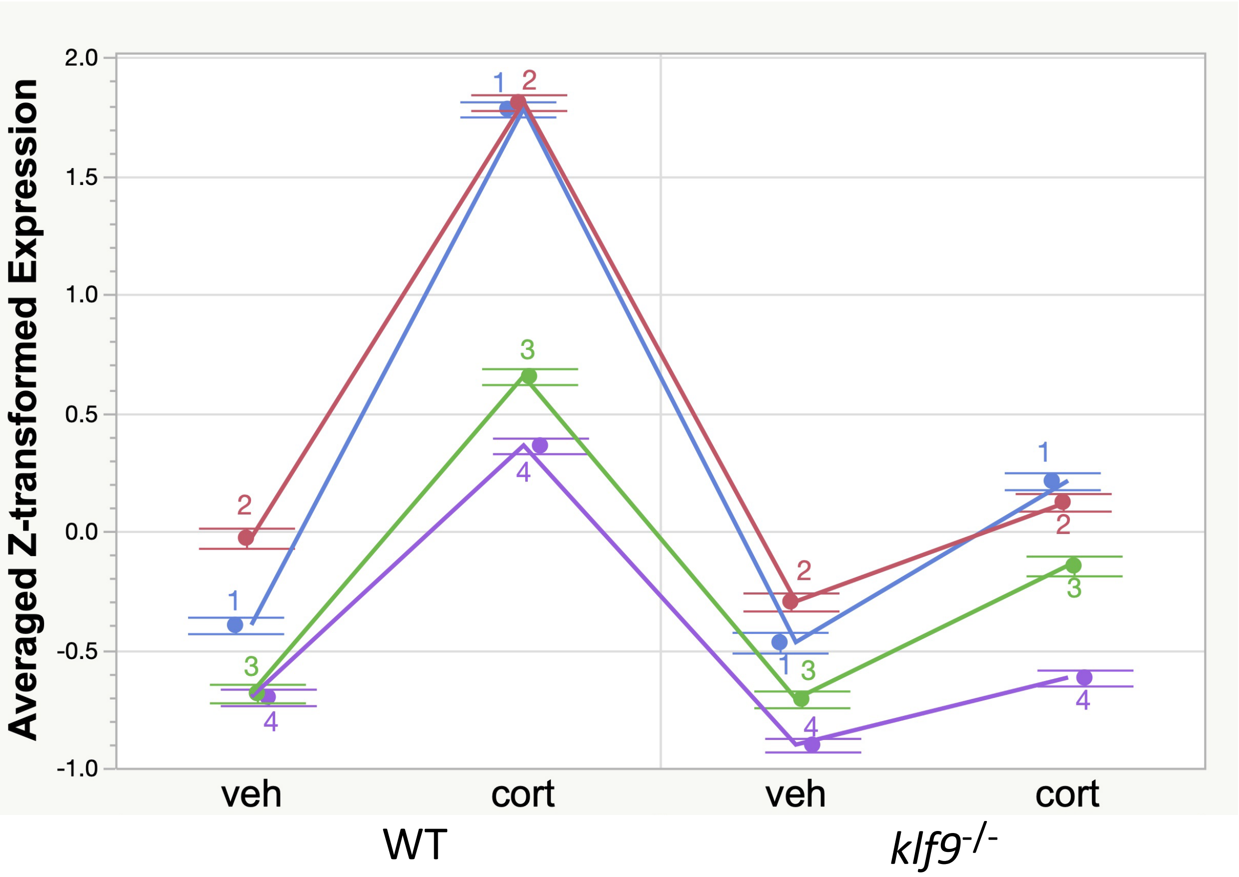


**Figure S12. Expression of 228 genes identified by hierarchical clustering to respond differently to cortisol in wild-type (WT) and klf9-/- larvae.** The expression data were rlog-transformed and then Z-normalized, and the plotted values are the average transformed expression across all genes under each condition in each of the four biological replicates, with error bars indicating the estimated standard error of the mean.

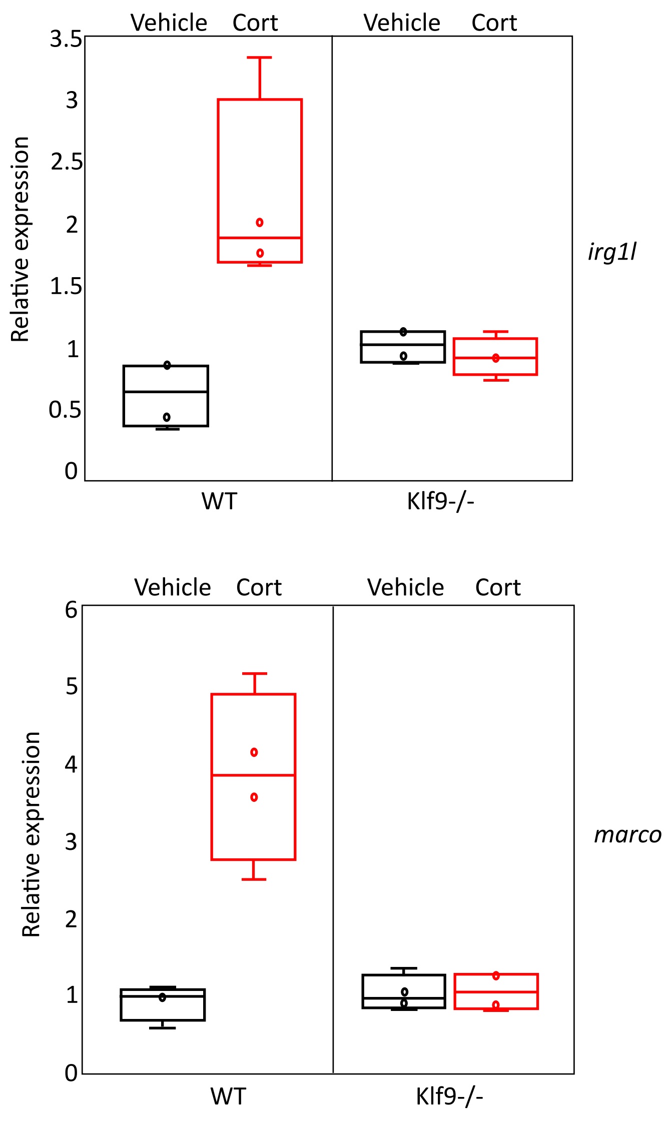


**Figure S13. Relative expression levels of *irg1l* and *marco* measured by qRT-PCR, in the same RNA samples that were subjected to RNA-seq** (see Fig. 4).

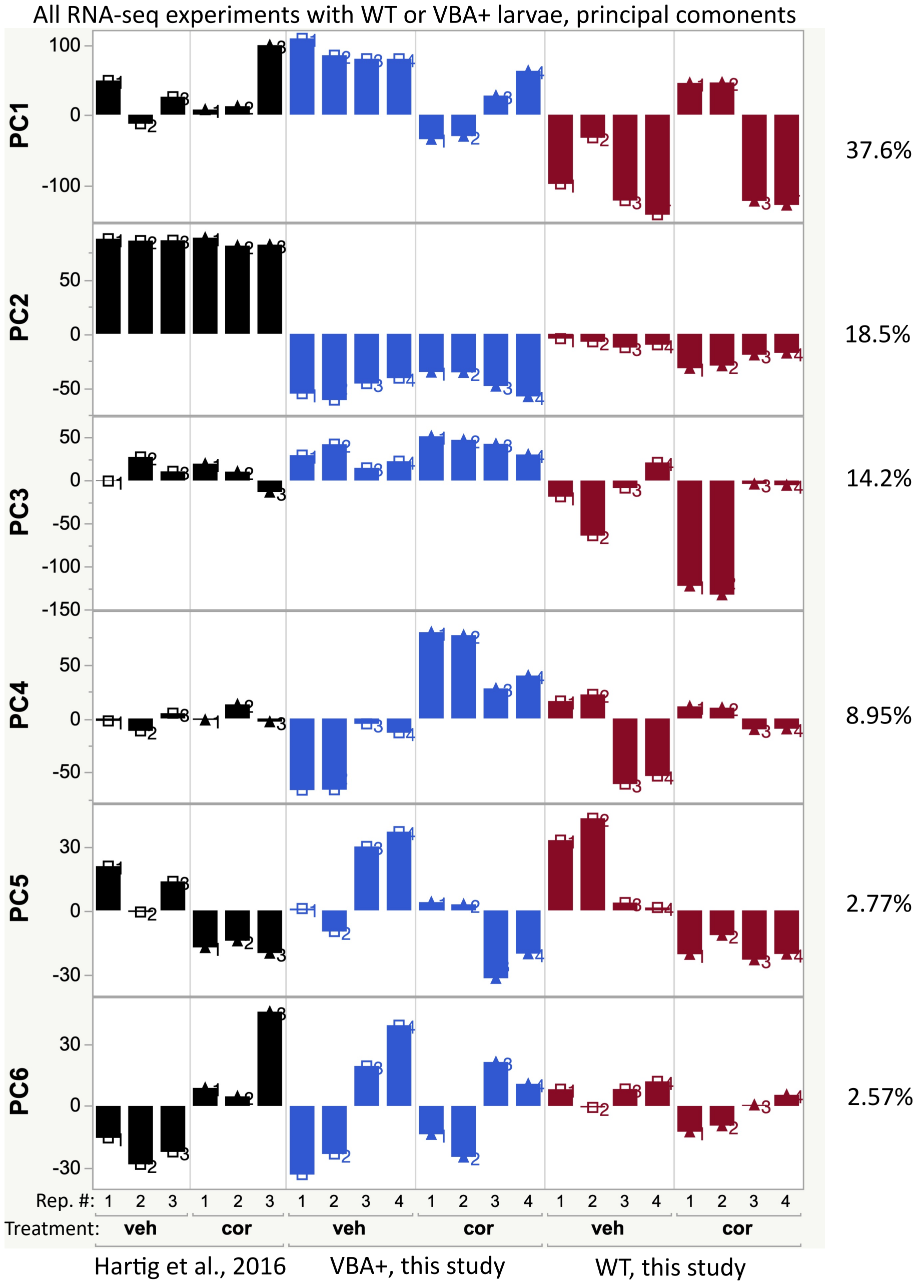


**Figure S14. Principal components of the variance in gene expression across three RNA-seq experiments examining the transcriptomic effects of chronic cortisol treatement.**
